# CASTER: Direct species tree inference from whole-genome alignments

**DOI:** 10.1101/2023.10.04.560884

**Authors:** Chao Zhang, Rasmus Nielsen, Siavash Mirarab

## Abstract

Genomes contain mosaics of discordant evolutionary histories, challenging the accurate inference of the tree of life. While genome-wide data are routinely used for discordance-aware phylogenomic analyses, due to modeling and scalability limitations, the current practice leaves out large chunks of the genomes. As more high-quality genomes become available, we urgently need discordance-aware methods to infer the tree directly from a multiple genome alignment. Here, we introduce CASTER, a site-based method that eliminates the need to predefine recombination-free loci. CASTER is statistically consistent under incomplete lineage sorting and is scalable to hundreds of mammalian whole genomes. We show both in simulations and on real data that CASTER is scalable and accurate and that its per-site scores can reveal interesting patterns of evolution across the genome.

---

The availability of genomes from a diverse array of species has rapidly increased (*1*), providing renewed hope for accurate species-level phylogenetic reconstruction (species trees) while modeling changes across the genome (gene trees) (*2*). However, the methodology for utilizing genome-wide data lags behind data availability. The traditional method of concatenating genes (*3*) is widely used and has reasonable accuracy in simulations (*4*) but fails to account for discordance across the genome due to Incomplete Lineage Sorting (ILS), leading to statistical bias (*5*). The practical alternatives that have emerged in the past decade are two-step methods that first reconstruct gene trees from demarcated loci and then combine them into species trees (*6*). This approach, widely used in practice (*7–9*), has computational advantages but also several limitations.

In theory, the locus-based approach requires identifying recombination-free loci that are fully unlinked and randomly sampled across the genome. While many analyses ignore this requirement (*10*) for practical reasons such as the difficulty of sequencing, homology detection, or alignment, analyses with access to genome-wide data increasingly attempt to fulfill these requirements. Setting aside the immense challenge of finding such loci (*11*), this approach cannot utilize the entire genome, as the requirement of having unlinked loci enforces the exclusion of some data. For instance, Chen et al (*8*) use 1kbp windows at least 50kbp distant from each other in their analysis, and similar levels of subsampling are used routinely (*12–16*). Additionally, the need to avoid recombination compels the use of short “c-genes” (*17*); when short loci are employed, gene tree estimation tends to exhibit high errors, which can propagate to the species tree (*9, 18–20*). Practitioners often struggle to balance the conflicting requirements to use longer loci and the need to avoid recombination within loci but maintain abundant recombination across loci; they often rely on arbitrary heuristic approaches, such as windowing.

Alternative methods, which infer the species tree directly from sequence data without estimating gene trees, exist but have not been widely adopted or found applicable. Some of these methods use Hidden Markov Models (HMMs) that explicitly model changes in tree topology along the sequence (*21, 22*) due to ILS. These methods are highly accurate but are limited to a small number of species because their computational time is quadratic in the number of possible tree topologies. Other site-based methods that account for ILS use statistics of site patterns (*23–26*). In principle, these methods can use the entire genome (without demarcating loci) and avoid the pitfalls of gene tree estimation errors. However, current site-based methods are less frequently used because they have not generally outperformed two-step methods in simulations (*27*), have limited statistical guarantees (e.g., when rates change across the species tree), do not provide a detailed view of heterogeneity across the genome, and are less scalable. For example, to apply SVDQuartets on 241 placental mammal genomes, Foley et al (*28*) excluded *>*99.9% of the aligned genome in their analyses. However, these limitations are not inherent to the concept of using site patterns.

In this paper, we introduce a new approach that uses site patterns to directly estimate a species tree. Our proposed method, called CASTER (Coalescence-aware Alignment-based Species Tree EstimatoR) is consistent under models of ILS, gives detailed information about histories of individual regions, is far more scalable than alternatives, and allows several forms of rate heterogeneity. Using extensive simulations, we show that CASTER, with high accuracy can estimate phylogenetic topologies for hundreds of full recombining genomes using *all aligned sites*. Furthermore, we apply CASTER to several existing genomic datasets, with a focus on the mammal dataset (*28*).

## CASTER: site-based species tree estimation

To enable the direct use of whole genome alignments for species tree inference, we developed a statistically consistent yet scalable site-based species tree inference method called CASTER.

Statistical consistency here is the guarantee of convergence to the correct tree with sufficiently long sequence data generated under an assumed model. As in several existing methods (*23, 29*), CASTER is built on the fact that for any four species (a quartet), the most likely gene tree under the multi-species coalescent (MSC), (*30*) and some other models of gene tree discordance (*31–33*), is the species tree. Examining each site (instead of inferred gene trees), CASTER assigns a score to each of the possible three quartet topologies for all quartets (Fig. 1A). Let *W* (*S, M*) be the score of a candidate species tree topology *S* with respect to the input alignment matrix *M* of length *L*. When *S* has only four species, we define,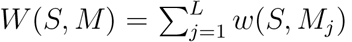 where *w*(*S, M*_*j*_) is the assigned weight of the site pattern in the *j*-th column of *M* . Further extending *S* to *n* species by summing over all 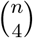quartets, we define

**Fig. 1:**
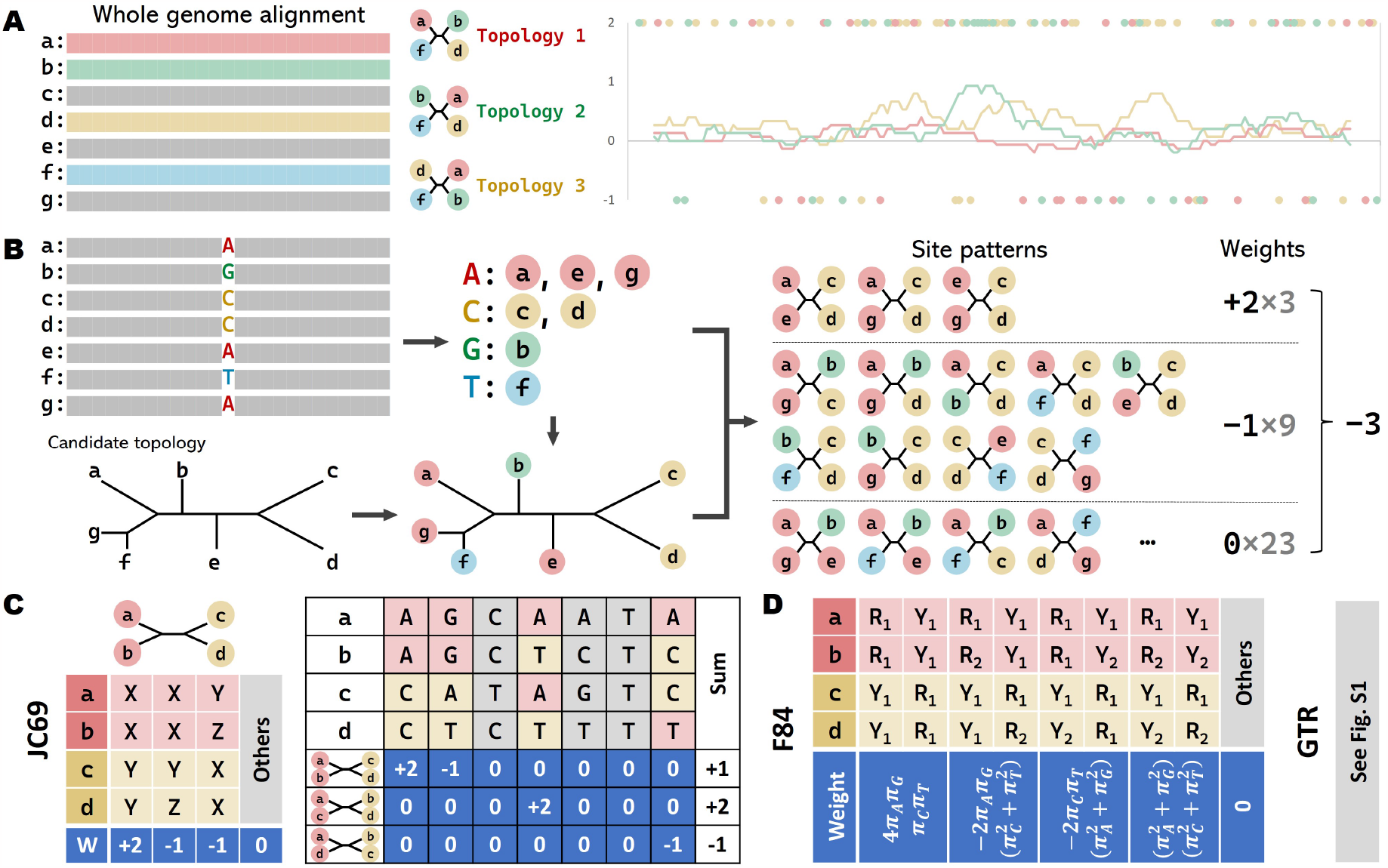
Overview of CASTER. (**A**) For each quartet of species, CASTER defines a weight, which can be positive, zero, or negative for each site pattern with respect to each of the three topologies; the weight of the topology is the sum of the site weights (shown as dots, zero weights omitted; lines: moving averages) across the whole genome. The species tree is the one with the highest weight. (**B**) For more than four leaves, to score a candidate tree, CASTER sums the weights of site patterns of all induced quartets. (**C**) Weights of site patterns for topology *ab cd* under the JC69 model and an example of counting weights of an MSA for all quartet topologies. *X, Y*, and *Z* represent distinct nucleotide letters. (**D**) Weights of site patterns for topology *ab cd* under the F84 model. *R*_1_ and *R*_2_ represent distinct purines, and *Y*_1_ and *Y*_2_ represent distinct pyrimidines. Similar matrix for GTR is shown in Fig. S1.

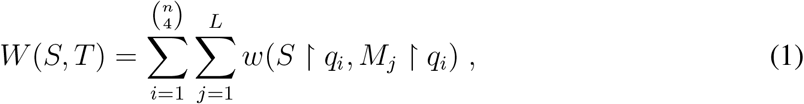

where *S* ∗ *q*_*i*_ and *M*_*j*_ ∗ *q*_*i*_ give the quartet subtree and site patterns for *i*-th quartet, *q*_*i*_. CASTER seeks *S* that maximizes *W* (*S, M*). The main contributions of CASTER are 1) defining three weighting schemes *w*(·, ·) for quartet site patterns that ensure statistical consistency, and 2) designing techniques that allow optimizing this score in a scalable fashion summing over all quartets *without* listing all quartets. We present outlines of each contribution, leaving a detailed treatment to materials and methods (*34*).

### Statistically consistent estimators from quartet site patterns

Each DNA site partitions taxa into up to four groups (Fig. 1B), and this division provides support for specific quartet topologies. For example, the site singled out in Fig. 1B provides support for putting species a and e together against c and d. In our attempt to quantify this support, we discovered that site patterns need to be weighted in surprising ways (Fig. 1C) to obtain a statistically consistent estimator of the true species tree topology (*34*) under the simple JC69 model (*35*). It gives us a way to score a site pattern for all quartet topologies (Fig. 1BC). Surprisingly, some patterns that seem intuitively to support (or at least not contradict) the species tree are given negative weights. These negative weights can be understood as correcting the bias caused by long-branch attraction (*36*).

To extend this approach to the F84 model (*22*), which allows variable equilibrium frequencies and different transition and transversion rates, we need a more sophisticated weighting scheme (Fig. 1C). The only required parameters are base frequencies, which can be trivially estimated. To further accommodate the General Time Reversible (GTR) model (*37*), which has three base frequency parameters and 6 rate parameters, we needed a different scheme that uses patterns across pairs of sites (Fig. S1). Using paired-site patterns, we extend the algorithm and its theoretical guarantees (*34*) to the GTR model. To do so, we need to partition sites to pairs and assume that for each pair of sites, they mutate independently but share the same evolutionary history, the same equilibrium frequencies, and the same substitution rate matrix.

### Scalable optimization

Maximizing the objective function defined in Equation (1) faces two challenges. Like any phylogenetic method, we need to contend with the large tree space. Additionally, for a given tree, computing Equation (1) would be slow if done naively, as iterating through all quartets by brute force would take Θ(*n*^4^) time. Existing methods, including SVDQuartets (*23*), rely on such brute-force approaches for small datasets or subsampling quartets. In contrast, we were able to design an algorithm to optimize the sum over *all* quartets without ever iterating through them. This approach colors the tips of a tree with the nucleotides and uses the resulting division into four groups to compute the total score in one calculation (Fig. 1B).

Having solved the problem of computing the score, we designed a search algorithm detailed in materials and methods inspired by weighted ASTRAL. Broadly, CASTER first uses the step-wise addition strategy to sequentially grow multiple trees, differentiated by the random order in which taxa are added. During step-wise additions, CASTER uses dynamic programming to guarantee each taxon is placed at its optimal position according to Equation (1). As a result, each of the multiple final trees is asymptotically optimal, regardless of the placement order. To achieve high accuracy on limited data, CASTER creates multiple such trees and synthesizes them into a final tree using dynamic programming, guaranteeing to find the optimal tree among all possible ways of combining them.

### Evaluation in Simulations

#### Recombination-based whole genome simulations

To capture the full complexity of molecular evolution, we simulated a dataset (SR201) of whole-genomes of 201 species (including one outgroup) under the Hudson coalescent model (*38*), which simulates recombination along the genome, allowing dependencies among local gene trees. To avoid unrealistic strict molecular clocks (i.e., ultrametricity), we varied the generation time and population size across the species tree. Similarly, we varied the mutation rate across the genome (*34*). We compared two versions of CASTER (CASTER-site for the F84 model and CASTER-pair for the GTR model) with three state-of-the-art methods that representing different phylogenetic approaches. We used the maximum-likelihood method RAxML-ng (*39*) with the concatenated alignment of the entire genome, which is statistically inconsistent (*5*) but widely used. We used the invariant-based site-based method SVDQuartets (*23*), also applied to the entire genome. We also included the two-step pipeline of gene tree inference using RAxML-ng and species tree summarization using wASTRAL (*40*) (RAxML-ng+wASTRAL shortened to wASTRAL henceforth). Consistent with common practice, to minimize recombination within loci, instead of giving the entire genome to wASTRAL, we gave it loci of size 500bp selected out of 2500bp windows, which accounts for only 20% of the data given to the other methods.

Both CASTER-site and CASTER-pair were more accurate and much faster than alternative methods (Fig. 2A and S2A). The mean error of CASTER-pair species tree errors was 1.3 branches compared to 1.7 branches for CASTER-site, which was a statistically significant improvement (see Table S1 for p-values of statistical tests substantiating all claims of significance throughout the paper). The mean errors were significantly higher for RAxML-ng (2.9 branches), SVDQuartets (3.8 branches), and wASTRAL (3.1 branches). CASTER-site and CASTER-pair took 4.5 and 34 core hours on average, compared to 282, 402, and 690 for RAxML-ng, SVDQuartets, and wASTRAL, respectively. Thus, CASTER-site is at least 60 times faster than alternative methods *and* more accurate.

**Fig. 2:**
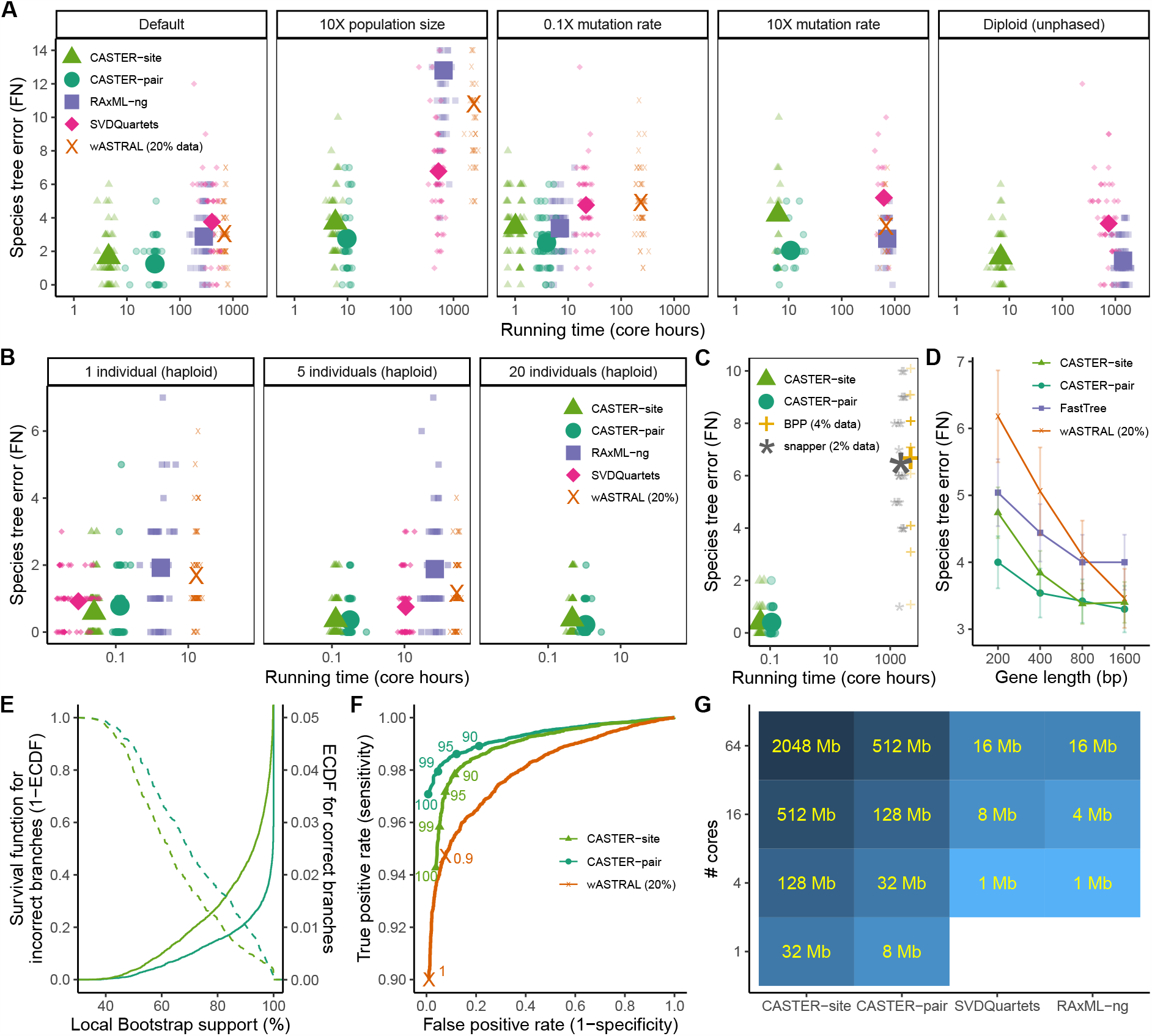
Benchmarking via simulations. The number of branches differing between inferred and true species trees (i.e., bipartition false negatives; FN) versus running time for five species tree inference methods under (**A**) various conditions of the SR201 dataset and (**B**) various numbers of individuals per species for the SR21 dataset. Small shapes denote the outcomes of each replicate; large and opaque shapes denote mean values. The y-axis is cut off at FN=14. (**C**) FN of BPP and snapper for two individuals per species on the SR21 dataset. To produce any results within a month of running time, we had to subsample 4% and 2% of sites for BPP and snapper. (**D**) Mean and standard error of FN versus gene length on the S101 dataset. (**E**) Empirical CDFs and (**F**) ROC curves of support values produced by CASTER on SR201 datasets summarized over 40,000 branches in total. (**G**) Scalability of various species tree inference methods versus the number of physical CPU cores used; we show the maximum number of sites (per each of 200 species) that can be analyzed within 3,000 core hours and 240 GB memory.

To investigate the effect of higher ILS, we increased the effective population size from 10^5^ to 10^6^. The increased ILS levels led to higher species tree estimation error levels for all the methods (Fig. 2A and S2B), but the level of impact varied across methods. Compared to the default condition, the advantages of CASTER over other methods became even more prominent; e.g., the error of RAxML-ng increased by 360% compared to a 120% increase for CASTER-pair. Interestingly, the consistent method wASTRAL also had significantly increased error (190%), perhaps due to the increasing levels of recombination within loci. Increasing population size retained or magnified the running time advantages of CASTER over other methods.

We varied the nucleotide substitution rate from 5 × 10^−8^ per site per generation to 5 × 10^−9^ and 5×10^−7^. Varying the mutation rate affected the accuracy and running time of all methods to varying degrees. Decreasing the mutation rate led to fewer substitutions and significantly lower accuracy (Fig. 2A) for all methods. Under this condition, CASTER-pair remained the most accurate method, followed by RAxML-ng, which had significantly higher error and 1.8 times longer running times. Note that lower mutation rates reduced running time for all methods (up to 19X). Increasing the substitution rate *also* increased error for CASTER-site and to a lesser degree for CASTER-pair and SVDQuartets (Fig. 2A); in contrast, the accuracy of RAxML-ng and wASTRAL remained essentially unchanged. It appears that methods based on quartet site patterns are more sensitive to high rates of evolution than those using maximum likelihood. Nevertheless, under the condition with higher rates, CASTER-pair was still more accurate than RAxML-ng and other methods.

CASTER computes branch supports using 1000 local block bootstraps by default at no additional computational cost beyond what is reported (*34*). Measuring the accuracy of these support values, we found that most incorrect branches had low support (Fig. 2E). Branches with 100% support in CASTER-site and CASTER-pair were correct 96.2% and 99.3% of the time, respectively, and 94.3% and 97.1% of correct branches had full support (Fig. 2E). Overall, the support values of CASTER were more predictive of accuracy than wASTRAL (Fig. 2F), the only other method that produces support by default and at no additional cost.

#### Multiple haplotypes or individuals per species

Unphased diploid genomes are commonly used in phylogenetic analyses. We tested the performance of CASTER-site, RAxML-ng (GTGTR4 model), and SVDQuartets using unphased genomes. CASTER-site and SVDQuartets can handle an unphased genome by arbitrarily phasing it and treating it as two individuals with haploid genomes; in contrast, RAxML-ng implements a model designed explicitly for diploid genomes (GTGTR4). Using unphased diploid genomes instead of haploid genomes resulted in a small but statistically insignificant improvement in the mean species tree errors for CASTER-site and SVDQuartets while RAxML-ng showed a significant improvement with a 50% decrease in error (Fig. 2A). RAxML-ng-GTGTR4 showed a small (12%) reduction in the error compared to CASTER-site, but this improvement came at the cost of a 200× higher running time.

To evaluate the performance of CASTER with multiple individuals per species, we also simulated a second dataset (SR21) of whole-genome alignments of 21 species (including one outgroup) with 20 haploid individuals per species (Fig. 2B). CASTER was easily scaled to 20 individuals, with running times never exceeding 3 hours. In contrast, SVDQuartets failed to run with 240 GB memory, and RAxML-ng and wASTRAL failed to finish within 3,000 core hours. Subsampling five individuals per species resulted in a significant degradation in accuracy for CASTER-pair (from 0.22 to 0.36 branches). With this reduced sampling, other methods also finished, but CASTER was the most accurate and scalable method. Further subsampling to one individual significantly reduced the accuracy of all methods (18% – 38% depending on the method), but CASTER remained the most accurate.

We also compared CASTER with two leading Bayesian approaches: the site-based method snapper (*25*) and the co-estimation method BPP (*41*). Since these two methods are very slow, to finish snapper and BPP runs in a reasonable time, we ran both methods, as well as CASTER, with two individuals per species. Furthermore, we had to subsample 2% of aligned sites for snapper and 4% of aligned sites for BPP. The species tree errors of snapper and BPP with sub-sampled sites (6.5 branches for both) were significantly higher than CASTER methods with all sites (0.4 branches for both) despite having 10,000X higher running time (Fig. 2C). When we also limited CASTER to the same input data as snapper and BPP, they were still substantially less accurate than CASTER (Fig. S2C). While the exact cause of the reduced accuracy of these methods, traditionally assumed to be very accurate, is unclear, we speculate that lack of ultrametricity (change of rates across the species tree) plays a role.

#### Recombination-free simulations

A surprising finding of our previous simulations was that wASTRAL was less accurate than the alternatives. A plausible reason is that wASTRAL was used with 5 times fewer data to avoid recombination, and that, despite this, recombination could still impact its results. To test this hypothesis, we compared CASTER to wASTRAL (with FastTree-II gene trees) and concatenation (using FastTree-II) on an existing (*42*) dataset (S101) of 101 species and 1000 recombination-free genes with various gene lengths. CASTER performed better than alternative methods regardless of gene length (Fig. 2D). With long genes (e.g., 1,600 bps), wASTRAL had similar error levels as CASTER-site and CASTER-pair, all of which were better than concatenation. With short genes (200 bps), however, CASTER-site (4.74) and CASTER-pair (4.0) were more accurate than both alternatives.

#### Scalability

Genome-scale phylogenomics requires the ability to handle larger amounts of data, making scalability a crucial factor. We simulated a concatenated genome alignment of 201 species and two billion base pairs to evaluate the scalability of four methods: CASTER-site, CASTER-pair, RAxML-ng, and SVDQuartets. The maximum amount of data that each method was able to handle within a 48-hour period was determined, given different computational resources in terms of CPU cores (1, 4, 16, and 64) and memory (4, 16, 64, and 240 GB). We inferred the species tree using the first 1, 2, 4, 8, …, and 2048 million base pairs (Mbps) and reported the maximum size each method could analyze. CASTER-site was able to consistently handle the largest amount of data (Fig. 2G), handling 32 Mbps with 1 physical core and 4 GB memory to 2048 Mbps (i.e., an entire mammalian genome) with 64 physical cores and 240 GB memory. The second most scalable method was CASTER-pair, which was also was able to handle up to 512 Mbps (a quarter of a mammalian genome) with 64 physical cores and 240 GB memory. RAxML-ng and SVDQuartets had *much* lower throughput compared to CASTER methods and were not able to handle more than 16 Mbps even with the maximum allowed resources. Thus, by handling significantly more data compared to other methods, CASTER may be able to improve the accuracy of genome-scale phylogenomics.We note that by using less computation on fixed dataset size, CASTER has the potential to reduce the carbon footprint of phylogenetic analyses.

### Applications to biological data

#### Mammals

We reanalyzed a recent dataset of 241 mammalian genomes (*28*). The main species tree published in the original analysis is inferred by SVDQuartets using 411,110 genome-wide nearly neutral sites. We reconstructed a species tree with CASTER-site using the aligned whole genomes, comprising 1.8 billion informative sites (Fig. S3A). Our whole-genome tree differs (Fig. 3A) from the published tree by two branches (among Afroinsectiphilia and Odd-nosed monkeys), but our topology is recovered in alternative analyses presented in the original publication. Running CASTER on 411,110 neutral sites used by Foley et al (*28*) revealed that these differences are not due to the choice of sites, but the choice of method (SVDQuartets vs CASTER). Remarkably, the entire analysis of 1.8 billion sites from 241 species (680 GB in total) took only 30 hours on a server node with 64 physical cores and 512 GB of allocated RAM.

**Fig. 3:**
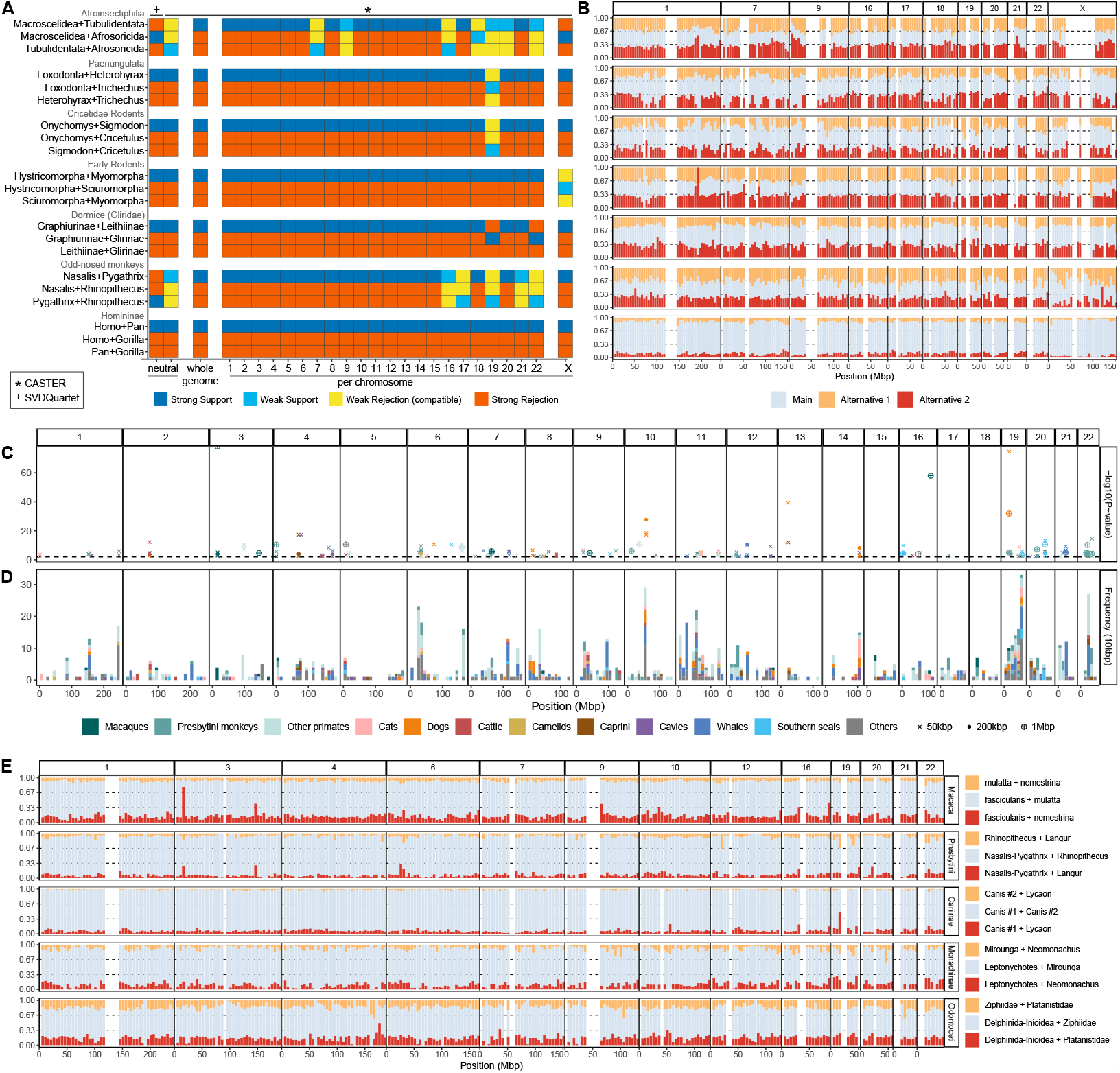
CASTER on the 241-taxon Mammalian dataset. (**A**) Species tree topologies inferred by CASTER-site using the whole genome, individual chromosomes, and 411,110 genome-wide nearly neutral sites are identical except for six branches shown here, in addition to the ancestor of Homininae, as an example of an easy branch. For each branch, we show which of the three possible topologies around it are recovered and whether the support is low (∗ 50%). We compare to the SVDQuartet tree from the original publication. (**B**) Normalized CASTER scores for main and alternative topologies of each branch featured in (A) averaged over 5Mbp sliding windows for select chromosomes. Regions missing from the alignment and low branch-specific alignment coverage (i.e., too many gaps pertaining to quartets around the branch) are left blank (*34*). (**C**) Log-transformed p-values for statistical tests detecting alternative topologies with unusually high scores in sliding windows of 50k, 200k, or 1Mbp size, showing only windows with p-value *<* 0.01 after Benjamini-Hochberg correction. (**D**) A histogram corresponding to the number of significant points with unusually high scores in sliding windows of 10kbp size, similar to part (C). (**E**) Similar to (B), showing 5Mbp sliding windows of normalized scores for select chromosomes and for branches with many statistically significant regions in part (C). See also Fig. S3.

The power of CASTER becomes more evident when examining the signal across the genome. We reconstructed species trees using individual chromosomes and found that the chromosome-specific trees differed from the main tree on six branches, including the two where CASTER disagrees with the Foley et al (*28*) tree (Fig. 3A). To further explore these disagreements, we computed a normalized sum of the CASTER scores (*34*) for the main topology found by CASTER and its two alternative topologies (rearrangements around the main topology) for each branch of the species tree. Visualizing these scores over 5Mbp sliding windows revealed chromosome-wide patterns that explain the difficulty of resolving these relationships (Fig. 3B). When examining a straightforward phylogenetic branch (e.g., humans/chimpanzees/gorillas), there is high support for the main topology, and the frequencies of the alternative topologies largely align with ILS expectations. Difficult branches diverge from this clean pattern in several ways.

One source of difficulty is rapid radiation. For example, the resolutions of Afroinsectiphilia and early rodents show normalized CASTER scores very close to 1*/*3, indicating potential rapid radiations (Fig. 3B). Small changes in the analysis pipeline (e.g. alignment, choice of sites, etc.) can lead to one topology or the other being favored. The difficulty is compounded by specific areas that show regional non-ILS-like patterns. Specifically, for the resolution of Afroinsectiphilia, certain regions in chromosomes 7, 9, 21, and X exhibit support for one of the two alternative topologies exceeding what one would anticipate under incomplete lineage sorting alone. CASTER enables users to detect such anomalies and further interrogate them.

One source of such anomalies is homology or modeling errors. For example, the alternative topologies of Paenungulata and Cricetidae display scores with substantial variation across 5Mb windows of the genomes, pointing to potential data errors or more complex evolutionary processes, such as selection. Conversely, we note that in most of the displayed branches, chromosome 19 exhibits a different pattern compared to other chromosomes. This discrepancy could be attributed to its exceptionally high gene content, which increases its susceptibility to misalignment and incorrect orthology detection due to the abundance of paralogous sequences. Beyond evidence of radiation and homology errors, we identified potential signs of hybridization around specific branches. For instance, in the resolutions of Dormice and Odd-nosed monkeys, the two alternative topologies consistently display divergent normalized CASTER scores across the genomic landscape (Fig. 3B). These observations suggest the possibility of introgression (*43*) and offer a plausible explanation for the longstanding debates surrounding their phylogenetic resolution (*44–47*).

To further probe introgression, we conducted a thorough genome scan using varied window sizes (10k, 50k, 200k, and 1Mbp). We identified 1101 windows in total from autosomal regions without clear alignment problems, where alternative topologies displayed significantly (*34*) higher scores than expected (Fig. 3CD). The elevated scores observed in these windows could not be solely explained by incomplete lineage sorting. A large proportion of these signals originated from higher primates (e.g. macaques and Presbytini monkeys), which have high taxon sampling. We also recovered abundant signals within Felidae (cats), Canidae (dogs), Bos (cattle), and Camelidae (camelids) species; all these groups have been known to undergo introgressions (*13, 14, 48–50*). In addition, we also discovered mutilple introgression signals within Caprini (sheep-goats), Cavia (cavies), Cetacea (whales), and Monachinae (Southern seals) with a less well-established history of reticulate evolution. These groups account for 80% of significant cases.

Visualizing scores averaged in 5Mbp sliding windows for five select branches with many regions with significant deviations showed interesting patterns (Fig. 3E). Within the Macaca (macaques), our whole-genome tree positioned *M. nemestrina* as the sister to *M. fascicularis* and *M. mulatta*. However, several 5Mbp sliding windows on chromosome 3 strongly support *M. mulatta* as the outgroup. This pattern also appears on chromosomes 9 and 16, though to a lesser degree. This finding is broadly in line with Fig. 3C and corroborates recent studies suggesting bidirectional gene flow between *M. nemestrina* and *M. fascicularis* (*51,52*). We also found similar patterns in Presbytini monkeys. Surprisingly, both branches share the windows on chromosome 3 with abnormally high supports for alternative topologies. Besides chromosome 3, we also observed significant deviations on chromosomes 6, 12, 19, and 20. Those signals, together with statistical tests (Fig. 3C) show a non-ILS source of discordance. We, therefore, hypothesize that the high scores for alternative topologies in Presbytini monkeys are due to introgression, consistent with the conclusions of an earlier study (*53*). We also observed similar patterns within Caninae (dogs), Odontoceti (toothed whales), and Monachinae (Southern seals), which suggests ancient reticulate evolution within those clades.

It is important not to automatically interpret any deviation from ILS as evidence of introgression, as alignment of assembly errors could potentially generate similar signals, particularly in regions with duplications. In fact, we observe several extreme values of the score in regions with high paralogy. A notable example is the strong signal that we detected between the *Gorilla* and *Pan* species within a pseudogene (LOC728688) paralogous to UHRF1. We also note that out of 1101 high-score windows, 928 are 1 kbp in size and a significant number of these windows coincide in their genomic positions, in different positions in the tree, suggesting alignment/assembly errors. For further illustration, refer to Fig. S4, which showcases instances of pronounced signals in 1kbp windows. These observations suggested that our approach could be harnessed to detect regions of interest to detect both errors (e.g., misalignment) and real biological insights (e.g., introgressive regions).

#### Ruminants

We reanalyzed a dataset of whole-genome orthologous sequences consisting of 373 million base pairs obtained from 51 ruminants (*8*). Using CASTER-site and CASTER-pair, we were able to infer the species tree in 58 and 257 core hours, respectively. Both methods produced the same inferred species tree topology, with full support on all branches (Fig. 4A). The inferred tree by CASTER differed from the published tree, supported by both ASTRAL and RAxML, by only one branch. In the published tree, sitatunga was the sister to mountain nyala and bongo, while CASTER united sitatunga and bongo. The relationship between these species is uncertain, and sitatunga is known to be able to cross-breed with close relatives, including bongo (*54*), making hybridization a potential explanation. Consistent with this explanation, we observed that across the genome, two topologies had similar support and both had higher support than the third topology (Fig. 4B), a pattern that is inconsistent with ILS but consistent with introgression (*43*).

**Fig. 4:**
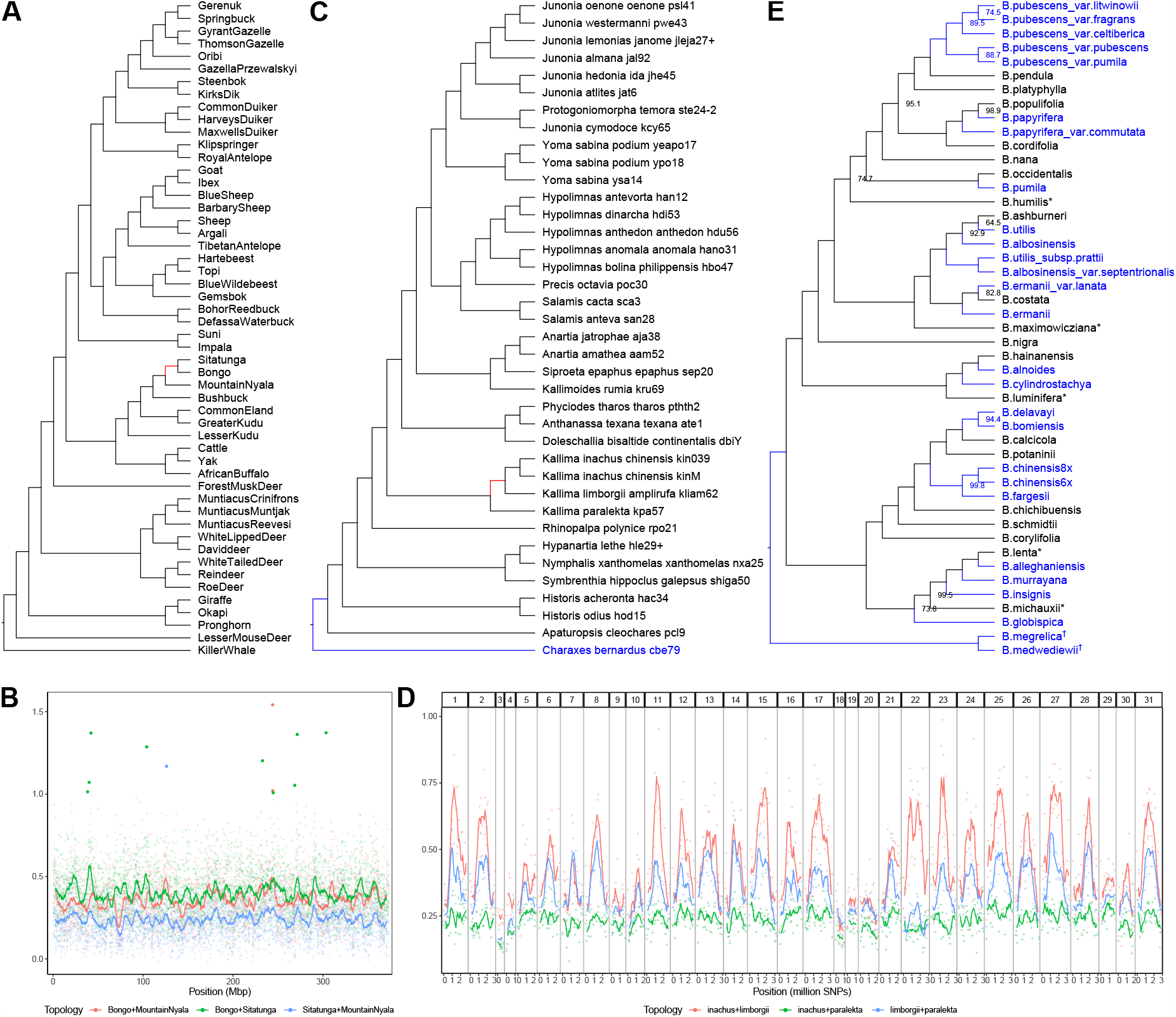
CASTER on the other real datasets. (**A**) CASTER tree based on 51 ruminant whole-genome orthologous sequences spanning 373 million base pairs. (**B**) CASTER-site scores for resolutions of *Sitatunga*/*Bongo*/*Mountain Nyala* across the aligned genomes. Lines represent 5Mbp moving averages, while dots depict 100kbp windows with extreme scores highlighted. (**C**) CASTER-site tree based on 38 whole-genomes of brush-footed butterflies, encompassing 166 million SNPs. (**D**) CASTER-site scores for resolutions of *Kallima* species across autosomes. Lines represent moving averages of 5M SNPs; dots depict 1M-SNP windows. (**E**) CASTER-pair tree based on 88,488 loci from 47 birch species. In (A), (C), and (E), branches colored red differ from the published trees, and branches colored blue are not present in the previously published trees; branches supports (%) are shown next to the branches, and 100% supports are omitted.

#### Butterflies

We reanalyzed a whole-genome sequencing dataset of 166 million SNPs from 38 brush-footed butterflies (Nymphalidae family), with a strong focus on the Nymphalinae subfamily (*55*). As only SNPs are available, we could only run CASTER-site for this dataset, which finished in 26 core hours. The inferred species tree topology by CASTER-site had full support on all branches and disagreed with the published analyses in a single branch (Fig. 4C). A small rearrangement was observed among Kallima (oak leaf butterflies). Unlike the published tree that put *K. paralekta* as a sister to *K. limborgii*, CASTER placed it as a sister to *K. limborgii* and *K. inachus*. The uneven scores of the two alternative topologies for this branch suggested by CASTER (Fig. 4D) indicate that there might be a significant amount of reticulate evolution among these species. Moreover, CASTER was able to place an outgroup (*Charaxes bernardus*) (a Satyrine) available in the alignment but not the published tree at the expected position relative to Nymphalines.

#### Lizards

We analyzed a dataset comprising 10,390 orthologous coding sequences, spanning a total length of 18 million base pairs, from 12 lizard species (*56*). Using both nucleotide and amino-acid sequences, we ran CASTER-site and CASTER-pair on multiple configurations, including with and without the third codon positions (Fig. S3B). All analyses yielded a fully supported species tree topology, consistent with both published ExaML and ASTRAL trees. Remarkably, all our runs finished in mere seconds on an eight-core desktop, demonstrating the efficiency and reliability of the CASTER algorithm.

#### Birch

We analyzed two RAD datasets of birch trees (*57*). A subset of data contains 50,970 loci from 20 diploid species and was analyzed by the original publication. On this dataset, the inferred tree by CASTER-pair matches the published tree by ASTRAL and by concatenation via RAxML (Fig. S3C). The inferred tree using CASTER-site differs from CASTER-pair by a single branch and matches the topology by ASTRID (Fig. S3D).

A second alignment also was available with 88,488 loci from 27 polyploid species (phased) and the 20 diploid species (Fig. 4E). This data was not used for species tree inference in the original publication due to polyploidy. The inferred tree by CASTER-pair on the dataset including polyploid species was compatible with the diploid tree. Notably, each polyploid species is placed as a sister to one of its putative diploid progenitor (e.g., *B. pumila*) or a clade of several putative diploid progenitors (e.g., *B. globispica*) except for *B. medwediewii* (a decaploid) and *B. megrelica* (a dodecaploid). *B. medwediewii* and *B. megrelica* are placed in a separate position as their putative diploid progenitors spread throughout the phylogenetic tree (*B. humilis*/*B. lenta*/*B. maximowicziana*/*B. michauxii*/*B. luminifera*).

### Discussion

While the use of genome-wide data for phylogenetic inference is routine, use of the entirety of alignable genomic data has hitherto been computationally prohibitive in most applications. CASTER provides a solution to this problem as the first consistent estimator of species phylogeny that is applicable to large whole-genome data sets . Our simulations demonstrate that CASTER has superior accuracy and scalability compared to other state-of-the-art methods. Furthermore, it is more robust, particularly in the CASTER-pair version, to high levels of ILS and rate variation. CASTER thus enables true whole-genome phylogenetic analyses even for large whole-genome data sets with many species and multiple individuals per species.

A main feature of CASTER is that it computes a score for every site. Interrogation of this score can help users identify the location of phylogenetic signal for various topologies across the genome. With a focus on mammals, we demonstrated CASTER’s potential in detecting genomic regions subject to introgression or assembly/alignment errors (Fig. 3C-E). Our main focus in the present work was species tree inference. However, we note that the per-site CASTER scores calculations could be further refined to provide a formalized framework for identifying introgression and distinguishing introgression signals from alignment/assembly errors. We also note that CASTER could potentially be improved by incorporating weighting by alignment quality into the objective function and by designing more accurate ways of pairing sites under the GTR model. Furthermore, the method could be extended to estimate branch lengths for the estimated species tree topology and could be adapted to accept various other types of data such as allele frequencies or binary characters. More ambitiously, future work may be able to extend the algorithm to the more challenging problems of inferring phylogenetic networks/admixture graphs.

## Acknowledgments

We thank Shuting Wang and Junyi Ding for their assistance in acquisition of published biological datasets. This work used Expanse at SDSC through allocation ASC150046 from the Advanced Cyberinfrastructure Coordination Ecosystem: Services & Support (ACCESS) program, which is supported by National Science Foundation grants.

## Data and materials availability

All data and code are available on Dryad (DOI: 10.5061/dryad.bg79cnph0). Up-to-date CASTER is available on GitHub (https://github.com/chaoszhang/ASTER).

