## Supplementary material for "CASTER: Direct species tree inference from whole-genome alignments": (34)

### Supplementary Materials for CASTER: Direct species tree inference from whole-genome alignments

### Materials and Methods

#### CASTER

##### Models and notations

CASTER imposes no constraints on the properties of the species tree. The species tree can have arbitrary population sizes and nucleotide substitution rates, including heterotachy, at different time points of any branch. Additionally, each site (under F84) or site pair (under GTR) can have its own evolutionary history or parameters. We describe the models below, referring the reader to Fig. S5 for an illustration.

**Species tree and sequence evolution model** The true species tree is represented by a binary tree $\mathbf{S} = (\mathbf{V_S}, \mathbf{E_S}, \mathbf{\Pi}, \mathbf{Q})$, where $\mathbf{V_S}$ denotes the set of internal and leaf nodes, and $\mathbf{E_S}$ denotes the set of branches including the root branch. Additionally, $\mathbf{\Pi} = (\Pi_1, \ldots, \Pi_k)$ represents the list of equilibrium frequencies for all $k$ genes, and $\mathbf{Q} = (Q_1, \ldots, Q_k)$ represents the list of substitution rate matrices for all genes. Note that here we use the term gene to refer to c-genes; i.e., an arbitrarily small region (e.g., one site) in the genome that shares the same evolutionary history. Each node $\mathbf{v} \in \mathbf{V_S}$ is a tuple of the species label $\sigma_\mathbf{v}$ and the date $\tau_\mathbf{v}$, denoted as $\mathbf{v} = (\sigma_\mathbf{v}, \tau_\mathbf{v})$. Here, $\sigma_\mathbf{v}$ is an extant species, and $\tau_\mathbf{v}$ denotes the date or age of $\mathbf{v}$. The leafset $\mathbf{L}$ comprises nodes $\mathbf{v}$ where $\tau_\mathbf{v} = 0$, and its size is denoted by $n = |\mathbf{L}|$. Each branch $\mathbf{e} \in \mathbf{E_S}$ is a tuple of the tail node $\mathbf{u_e}$, head node $\mathbf{v_e}$, and functions $\mathbf{N_e}$ and $\mathbf{M_e}$. The function $\mathbf{N_e} : [\tau_{\mathbf{v_e}}, \tau_{\mathbf{u_e}}) \to \mathcal{R}_+$ maps a time point $t$ on branch $\mathbf{e}$ to the effective population size at time $t$. Furthermore, the list $\mathbf{M_e} = (\mu^1_\mathbf{e}, \ldots, \mu^k_\mathbf{e})$ contains functions where each $\mu^i_\mathbf{e} : [\tau_{\mathbf{v_e}}, \tau_{\mathbf{u_e}}) \to \mathcal{R}_+$ maps a time point $t$ on branch $\mathbf{e}$ to the nucleotide substitution rate of the $i$-th gene at time $t$. For the root branch, we let $\tau_{\mathbf{u_e}} = +\infty$ by convention.

**Gene trees** The set of gene trees $\mathcal{G} = \mathbf{G}_1, \ldots, \mathbf{G}_k$ is generated independently at random from the true species tree $\mathbf{S}$ under the multi-species coalescent (MSC) process (*1*). Each gene tree $\mathbf{G}_i$ is represented by a tuple $(\mathbf{V}_i, \mathbf{E}_i, \Pi_i, Q_i)$, in which $\mathbf{L} \subseteq \mathbf{V}_i$ . For each branch $\mathbf{e} \in \mathbf{E}_i$ of gene tree $\mathbf{G}_i$ at a time $t \in [\tau_{\mathbf{v_e}}, \tau_{\mathbf{u_e}})$, we can find a corresponding branch $\mathbf{e}' \in \mathbf{E_S}$ in the true species tree such that $\mathbf{N_e}(t) = \mathbf{N_{e'}}(t)$ and $\mu_{\mathbf{e}}(t) = \mu^i_{\mathbf{e}'}(t)$.

**Sites or site pairs** Under the site model (JC69 or F84), a single nucleotide $\mathbf{p}_i$ is randomly evolved along the branches of each gene tree $\mathbf{G}_i$. As we will discuss in Practical measures, the restriction to a single site is not necessary and is used only for ease of exposition. At the root of $\mathbf{G}_i$, $\mathbf{p}_i$ is sampled from the set $\Sigma = \{A, C, G, T\}$ with frequencies $\Pi_i$. At any time point $t$ along a gene tree branch $\mathbf{e}$, the substitution rate matrix of $\mathbf{p}_i$ is $\mu_{\mathbf{e}}(t)Q_i$. For convenience, we use $p^i_{\mathbf{v}}$ or $p^i_{\sigma_{\mathbf{v}}}$ to denote the nucleotide $\mathbf{p}_i$ at node $\mathbf{v}$. Under the pair model (GTR), a pair of nucleotides are randomly and independently evolved along each gene tree $\mathbf{G}_i$. The two nucleotides share the same equilibrium frequencies $\Pi_i$ and the same substitution rate matrix $\mu_{\mathbf{e}}(t)Q_i$ at each time point of each branch. In this case, we use $\mathbf{p}_i$ to denote the pair of sites, and let $p^i_{\mathbf{v}}$ or $p^i_{\sigma_{\mathbf{v}}}$ denote the pair of nucleotides $\mathbf{p}_i$ at node $\mathbf{v}$.

##### Objective function

We next reformulate Equation (1) more formally. For any species tree topology $S_*$, let $\mathbf{T}(S_*) = \{T_1, \ldots, T_{\binom{n}{4}}\}$ be the set of all quartet tree topologies in $S_*$. CASTER looks for the optimal species tree topology $S_*$ by maximizing the following objective function:

$$W(S_*) = \sum_{T \in \mathbf{T}(S_*)} \sum_{i=1}^{k} w_i(T) \tag{2}$$

where $w_i(T)$ depends on the model. For every site $\mathbf{p}_i$ and every four species $a, b, c, d$, we let $w_i(ab|cd)$ denote the weight of the site pattern $p^i_a p^i_b | p^i_c p^i_d$. Weights for JC69 and F84 models are defined in Fig. 1CD and weights for GTR are defined in Fig. S1.

**Theorem 1.** *Maximizing $W(S_*)$ in Equation (2) creates a statistically consistent estimator for the topology of the true species tree* $\mathbf{S}$ *under JC69, F84, or GTR models using the associated site pattern weights.*

Proof of Theorem 1 and the following theorems are provided in the proofs section of the supplement. To summarize, the proof first shows that the site patterns and their weights are carefully engineered so that for a species tree $\mathbf{S}$ with only four leaves and for every single gene, the expected weight of the topology matching $\mathbf{S}$ is greater than the two alternatives in expectation. More precisely, when the earliest gene tree coalescence is not a deep coalescence, the gene tree topology must match $\mathbf{S}$; in this case, the site patterns have a higher expected weight for $\mathbf{S}$ than the two alternatives; When there is deep coalescence and an opportunity for topological discordance, by symmetry, the expected scores of all three topologies are the same. Thus, $\mathbf{S}$ maximizes the expected score of every gene. Since all gene trees are independently sampled from $\mathbf{S}$, then maximizing $W(S_*)$ in Equation (2) creates a statistically consistent estimator for $\mathbf{S}$. This conclusion can be then extended to species trees with more than four species by summing over the species tree restricted to all quartets of species.

##### Optimization algorithm

**Overview** At a high level, CASTER infers the species tree using two steps. The first step follows a greedy search heuristic:

1. Start with a backbone tree of three randomly selected species.

2. At each step, select an unplaced species $a$ and place it onto the backbone tree $S_*$ using Algorithm 1 on its optimal position. To obtain the optimal placement, we first need to compute total weights for all tripartitions for all possible placements of $a$. There are only $O(n)$ such tripartitions, which all fall into the following categories:

(a) $(P_1 \cup \{a\}|P_2|P_3)$ or $(P_1|P_2 \cup \{a\}|P_3)$ or $(P_1|P_2|P_3 \cup \{a\})$

(b) $(P_1 \cup P_2|\{a\}|P_3)$ or $(P_1|\{a\}|P_2 \cup P_3)$.

where $(P_1|P_2|P_3) \in \mathcal{P}(S_*)$. In Algorithm 1, we compute the total weights for all tripartitions in $O(nk)$ time using a post-order traversal.

3. Repeat step 2 until all $n$ species are placed on the backbone tree.

This optimization algorithm guarantees the reconstruction of the true species tree topology provided with sufficient data.

**Theorem 2.** *The result of the greedy search used in CASTER is a statistically consistent estimator for the topology of the true species tree* $\mathbf{S}$.

The time complexity of Algorithm 1 is $O(n^2k)$ and memory consumption is linear making it a very scalable method. To achieve the $O(n^2k)$ running time, CASTER avoids listing all quartets. Instead, it uses the notion of scoring tripartitions, as detailed below. Further optimizations described in Practical measures improve the memory consumption at the cost of a slight increase in time complexity to $O(n^2k \log n)$.

With infinite sites, the greedy algorithm produces the same topology regardless of the order of placing species. However, with finite sites, placing species in different orders can produce different and suboptimal topologies. To improve optimization, we run the greedy algorithm multiple times in different orders to obtain a set of trees. We then use Algorithm 2 to find the optimal tree topology constrained to the tripartitions in the greedy trees.

**Tripartition scoring** To avoid listing all quartets, CASTER uses an algorithmic trick inspired by (weighted) ASTRAL (*2, 3*). Noting that an internal node in an unrooted tree divides taxa into three groups (a tripartition), CASTER uses the total weights of all quartets constrained by

**Algorithm 1** The greedy placement algorithm. For each node $\mathbf{v}$, let $\mathbf{L_v}$ denote the leafset under the subtree of $\mathbf{v}$. For each leaf node $\mathbf{v}$, let $\mathbf{1_v}$ denote the one-hot matrix for nucleotides of $\mathbf{v}$. For example, under the JC69 model, if $p^i_{\mathbf{v}} = A$, then $\mathbf{1_v}[i, A] = 1, \mathbf{1_v}[i, C] = \mathbf{1_v}[i, G] = \mathbf{1_v}[i, T] = 0$. Let $\mathbf{L}_{S_*}$ denote the leafset of $S_*$.

**Require:** $W(P_1|P_2|P_3)$ as defined in Equation 4.

**Ensure:** $W$ maps tripartitions for potential placements to their total weights

- **procedure** GREEDYALGORITHM
  - $L_0 \leftarrow$ three randomly chosen taxa from $\mathbf{L}$
  - $S_* \leftarrow$ An arbitrary tree with leafset $L_0$ ▷ The starting tree
  - **for** $a \in \text{shuffle}(\mathbf{L} - L_0)$ **do** ▷ Placing $a$ onto $S_*$
    - Reroot $S_*$ at an arbitrary leaf
    - $W \leftarrow$ an empty look-up table
    - $\mathbf{\Sigma} \leftarrow \sum_{\mathbf{v} \in \mathbf{L}_{S_*}} \mathbf{1_v}$
    - WEIGHTRECURSION(the heavy child of the root of $S_*, W, \mathbf{\Sigma}$)
    - $S_* \leftarrow$ the optimal tree by OPTIMALTOPOLOGY($W$) ▷ See Alg. 2
  - **return** $S_*$
- **procedure** WEIGHTRECURSION($\mathbf{w}, W, \mathbf{\Sigma}$)
  - **if** $\mathbf{w}$ is leaf node **then**
    - $W[\mathbf{L_w}|\{a\}|\mathbf{L}_{S_*} - \mathbf{L_w}] \leftarrow W(\mathbf{L_w}|\{a\}|\mathbf{L}_{S_*} - \mathbf{L_w})$
      ▷ $O(k)$ time using $\mathbf{\Sigma}_1 = \mathbf{1_w}$, $\mathbf{\Sigma}_2 = \mathbf{1}_a$, $\mathbf{\Sigma}_3 = \mathbf{\Sigma} - \mathbf{1_w}$ in (4),(6)
    - **return** $\mathbf{1_w}$
  - **else**
    - $(\mathbf{u}, \mathbf{v}) \leftarrow$ the children of $w$ with higher and lower subtree height
    - $\mathbf{\Sigma_u} \leftarrow$ WEIGHTRECURSION($\mathbf{u}, W, \mathbf{\Sigma}$)
    - $\mathbf{\Sigma_v} \leftarrow$ WEIGHTRECURSION($\mathbf{v}, W, \mathbf{\Sigma}$)
    - $W[\mathbf{L_w}|\{a\}|\mathbf{L}_{S_*} - \mathbf{L_w}] \leftarrow W(\mathbf{L_w}|\{a\}|\mathbf{L}_{S_*} - \mathbf{L_w})$
      ▷ use $\mathbf{\Sigma}_1 = \mathbf{\Sigma_u} + \mathbf{\Sigma_v}$, $\mathbf{\Sigma}_2 = \mathbf{1}_a$, $\mathbf{\Sigma}_3 = \mathbf{\Sigma} - (\mathbf{\Sigma_u} + \mathbf{\Sigma_v})$ in (4),(6)
    - $W[\mathbf{L_u} \cup \{a\}|\mathbf{L_v}|\mathbf{L}_{S_*} - \mathbf{L_w}] \leftarrow W(\mathbf{L_u} \cup \{a\}|\mathbf{L_v}|\mathbf{L}_{S_*} - \mathbf{L_w})$
      ▷ use $\mathbf{\Sigma_u} + \mathbf{1}_a$, $\mathbf{\Sigma_v}$, and $\mathbf{\Sigma} - (\mathbf{\Sigma_u} + \mathbf{\Sigma_v})$
    - $W[\mathbf{L_u}|\mathbf{L_v} \cup \{a\}|\mathbf{L}_{S_*} - \mathbf{L_w}] \leftarrow W(\mathbf{L_u}|\mathbf{L_v} \cup \{a\}|\mathbf{L}_{S_*} - \mathbf{L_w})$
      ▷ use $\mathbf{\Sigma_u}$, $\mathbf{\Sigma_v} + \mathbf{1}_a$, and $\mathbf{\Sigma} - (\mathbf{\Sigma_u} + \mathbf{\Sigma_v})$
    - $W[\mathbf{L_u}|\mathbf{L_v}|\{a\} \cup \mathbf{L}_{S_*} - \mathbf{L_w}] \leftarrow W(\mathbf{L_u}|\mathbf{L_v}|\{a\} \cup \mathbf{L}_{S_*} - \mathbf{L_w})$
      ▷ use $\mathbf{\Sigma_u}$, $\mathbf{\Sigma_v}$, and $\mathbf{1}_a + \mathbf{\Sigma} - (\mathbf{\Sigma_u} + \mathbf{\Sigma_v})$
    - **return** $\mathbf{\Sigma_u} + \mathbf{\Sigma_v}$

**Algorithm 2** Computing the optimal tree topology $S_*$ maximizing $W(S_*)$ constrained to all tripartitions in $W[\cdot]$. For simplicity, we only show how to compute the maximum $W(S_*)$, and the optimal $S_*$ can be trivially derived. Note that this Algorithm is run after Algorithm 1, which saves the weights in $W[\cdot]$. In the end, backtracking through partitions chosen in Line 13 gives the optimal tree.

**Require:** $W$ maps tripartitions in placed trees to their total weights
1: **procedure** OptimalTopology($W$)
2:     $a \leftarrow$ an arbitrary leaf species in $S_*$
3:     $M \leftarrow \emptyset$ ▷ for memoization
4:     OptimalTopologyDP($\mathbf{L}_{S_*} - \{a\}, W, M$)
5: **procedure** OptimalTopologyDP($P, W, M$)
6:     **if** $|P| = 1$ **then**
7:         $M[P] = 0$
8:     **else if** $P$ does not exist in $M$ **then**
9:         $M[P] \leftarrow -\infty$
10:         **for** each tripartition in $M$ in the form of $(P_1|P_2|\mathbf{L}_{S_*} - P)$ **do**
11:             OptimalTopologyDP($P - P_1, W, M$)
12:             OptimalTopologyDP($P - P_2, W, M$)
13:             $M[P] \leftarrow \max(M[P], W[P_1|P_2|\mathbf{L}_{S_*} - P] + M[P - P_1] + M[P - P_2])$

a tripartition as a building block to compute the total weights. For a tripartition $(P_1|P_2|P_3)$ and a gene $\mathbf{G}_i$, we define

$$W_i(P_1; P_2, P_3) = \sum_{a,d \in P_1} \sum_{b \in P_2} \sum_{c \in P_3} w_i(ad|bc) \tag{3}$$

where $w_i(ad|bc)$ is the weight of quartet $ad|bc$ in gene $\mathbf{G}_i$, and we similarly define $W_i(P_2; P_3, P_1)$ and $W_i(P_3; P_1, P_2)$; then,

$$W(P_1|P_2|P_3) = \sum_{i=1}^{k} W_i(P_1; P_2, P_3) + W_i(P_2; P_3, P_1) + W_i(P_3; P_1, P_2)\ . \tag{4}$$

Let $\mathcal{P}(S_*)$ denote the set of tripartitions corresponding to all internal nodes of a species tree topology $S_*$. Since counting all quartets constrained to elements of $\mathcal{P}(S_*)$ covers $\mathbf{T}(S_*)$ exactly twice, we have

$$W(S_*) = \frac{1}{2} \sum_{(P_1|P_2|P_3) \in \mathcal{P}(S_*)} W(P_1|P_2|P_3)\ , \tag{5}$$

where $W(S_*)$ is the score of the species tree topology $S_*$.

Crucially, the sums from (3) can be computed in a fast way without actually performing the sums using simple preprocessing. It is easiest to see how this can be done under the JC69 model. Let $\mathbf{G}_i$ be a gene and $(P_1|P_2|P_3)$ be a tripartition. We define $A^i_j = |\{\mathbf{v} : \mathbf{v} \in P_j, p^i_{\mathbf{v}} = A\}|$, $C^i_j = |\{\mathbf{v} : \mathbf{v} \in P_j, p^i_{\mathbf{v}} = C\}|$, $G^i_j = |\{\mathbf{v} : \mathbf{v} \in P_j, p^i_{\mathbf{v}} = G\}|$, and $T^i_j = |\{\mathbf{v} : \mathbf{v} \in P_j, p^i_{\mathbf{v}} = T\}|$, which represent the number of species in $P_j$ having nucleotide A, C, G, and T, respectively, at site $p_j$. We represent $A^i_j$, $C^i_j$, $G^i_j$, and $T^i_j$ as three matrices $\mathbf{\Sigma}_j[i, X] = X^i_j$ for $X \in \Sigma$. Given $\mathbf{\Sigma}_1, \mathbf{\Sigma}_2, \mathbf{\Sigma}_3$, we can compute Equation (3) in $O(1)$ time using site patterns and weights in Figure 1c, as

$$
\begin{aligned}
W_i(P_1; P_2, P_3) = +2 \times \sum_{X \in \Sigma} \sum_{Y \in \Sigma \setminus \{X\}} & \binom{\mathbf{\Sigma}_1[i, X]}{2} \mathbf{\Sigma}_2[i, Y] \mathbf{\Sigma}_3[i, Y] \\
-1 \times \sum_{X \in \Sigma} \sum_{Y \in \Sigma \setminus \{X\}} \sum_{Z \in \Sigma \setminus \{X, Y\}} & \binom{\mathbf{\Sigma}_1[i, X]}{2} \mathbf{\Sigma}_2[i, Y] \mathbf{\Sigma}_3[i, Z] \\
-1 \times \sum_{X \in \Sigma} \sum_{\{Y, Z\} \subseteq \Sigma \setminus \{X\}} & \mathbf{\Sigma}_1[i, Y] \mathbf{\Sigma}_1[i, Z] \mathbf{\Sigma}_2[i, X] \mathbf{\Sigma}_3[i, X] \ .
\end{aligned}
\tag{6}
$$

The same idea can extend Equation (6) to F84 and GTR models using more complex sets of counters, as explained in detail in the supplementary equations.

##### Practical measures

The following measures are employed to enhance sample complexity, running time, and memory consumption while maintaining the theoretical guarantees.

*Independence assumptions:* CASTER assumes known equilibrium frequencies per site. In practice, equilibrium frequencies are estimated for windows of the genome (10,000-bp intervals by default) and assumed to be fixed in each window. Moreover, while the model assumes independent evolution of one site (or pair) per gene, we use all sites of the aligned genomes in practice. Thus, we allow multiple sites per genealogy (and allow missing data). Because the expected score of any site is higher for the species tree compared to the alternative topologies, sampling multiple sites does not break consistency, as long as the number of sites sampled per genealogy is not adversarially designed to overrepresent topologies not matching the species tree. In fact, it can be proved that CASTER is statistically consistent under the Hudson coalescent model (*4*). We omit a formal proof, which can follow a simple idea. We can divide $k$ sites into $\lfloor\sqrt{k}\rfloor$ bins in a round-robin manner; i.e., $j$-th bin includes $i\lfloor\sqrt{k}\rfloor + j$-th sites ($0 < i \leq \lfloor\sqrt{k}\rfloor$). As $k \rightarrow +\infty$, sites in each bin are at linkage equilibrium (when $\lfloor\sqrt{k}\rfloor \rightarrow +\infty$) and therefore follow our assumed model. Thus, maximizing Equation 2 with respect to sites in any bin creates a statistically consistent estimator of the true species tree topology ($k/\lfloor\sqrt{k}\rfloor \rightarrow +\infty$). It follows that maximizing Equation 2 with respect to sites in all bins is also statistically consistent.

*Site pairing:* CASTER-pair is based on pairs of sites, which requires site pairing when applied to alignments. We first group sites by the number of unique nucleotide letters. In each group, we pair neighboring sites. To reduce the effect of recombination within pairs, we only keep pairs of sites that are less than 20 nucleotides apart in the original alignments.

*Multiple individuals:* CASTER can easily adapt to multiple haplotypes by summing over all haplotypes of each species in Equation (2). This approach allows the use of multi-individual datasets, representing each individual as one phased haplotype or two unphased haplotypes but with heterozygous positions masked for CASTER-pair. CASTER-site can also further utilize unphased sequences without masking heterozygous positions since it treats all sites as independent.

*Other data types:* CASTER-site can naturally take concatenated SNPs as inputs, with two caveats. Since invariable sites are not observed, CASTER would have to assume that equilibrium frequencies do not change dramatically across SNPs. Currently, CASTER-site does not natively support inputs in VCF/BCF formats. Instead, inputs should be first converted to

FASTA/PHYLIP formats as supermatrices.

CASTER-pair's nucleotide GTR model can be easily adapted to the amino acid GTR model. Patterns in Fig. S1 suggest that any partitioning of the 20 amino acid letters can be used to replace R/Y, M/K, and W/S to achieve statistical consistency under the amino acid GTR model. Currently, CASTER-pair supports amino acid sequences by applying the following trick: replacing amino acid letters C, M, I, L, and V in the input file with nucleotide letter A; D, E, Q, N, H, R, K with T; S, T, A, G, P as C; W, Y, F with G. Note that while any grouping works, we chose this grouping to match amino acid similarity according to BLOSUM matrix. We demonstrated the use of this treatment in the lizard dataset.

*Further scalability optimizations*: Although Algorithm 1 has an acceptable time complexity of $O(n^2k)$ and is easy to understand, it has a major shortcoming for handling truly genome-wide data: It requires memory allocation for storing the matrix $\mathbf{\Sigma_u}$ in each layer of the recursion, which can lead to a stack of size $O(k \log n)$. To reduce the memory consumption, we avoid saving $\mathbf{\Sigma_u}$ and $\mathbf{\Sigma_v}$ onto the stack; instead, we swap the order of lines 18 and 19, and recompute $\mathbf{\Sigma_v}$ by simply re-counting site patterns. Thus, we reduce the memory to $O(k)$ at the cost of slightly increasing the time complexity to $O(n^2 k \log n)$. Because this time complexity can be problematic for large $n$, we further reduce the running time for $n \geq 200$, using weighted ASTRAL's divide-and-conquer (i.e., two-step) approach. Instead of placing taxon one at a time, we first build a guide tree with a small subset of taxa and divide the rest of the taxa into clusters by placing them on this tree. Then, taxa in each cluster are added sequentially onto a separate copy of the guide tree, and the inferred species tree is then obtained by merging all guide trees without any extra computation. By making subsets limited to size $O(\sqrt{n})$ in expectation, we guarantee that this approach terminates in $O(n^{1.5+\epsilon}k)$ time.

##### Branch support

We compute branch support using local block bootstrapping. In each bootstrap replicate (default: 1000 replicates), we first divide the whole alignment into blocks of approximately equal sizes (default: 10,000 bps) and then randomly sample blocks with replacement such that the total length of sampled blocks approximately equals the length of the whole alignment. We compute support *locally* around each branch. For each internal branch in the inferred species tree and each bootstrap replicate, we compute the score of the quadrapartition defined by the internal branch and the scores of the two alternative topologies (NNI rearrangements). The branch support of each internal branch is defined as the number of bootstrap replicates where the score of the topology of the internal branch is higher than the scores of the alternatives.

#### Simulations

##### msprime simulations

We simulate aligned genomes with recombination using `msprime` (*5*); detalied parameters are listed in Tables S2 and S3. For the species tree (called the demographic model in msprime), we use species trees simulated in earlier work (*6*). These species trees, which include 200 ingroup and one outgroup species, were simulated under the Yule process using `Simphy` (*7*) with the tree height set to $10^7$ generations and the speciation rate set to $10^{-7}$ per generation. We divide all species tree branch lengths by 10, resulting in species trees with $10^6$ generations and a speciation rate of $10^{-6}$ per generation. To avoid ultrametricity, we then multiply each branch length in generations by a multiplier individually drawn from a Gamma distribution with mean $= 1$. The population size of each branch is independently drawn from a log-normal distribution with mean $10^5$ or $10^6$.

We simulate ancestries of diploid genomes using the Hudson model (*4*). We set the length to $5 \times 10^6$ bps and the recombination rate to $5 \times 10^{-8}$ per bp per generation. We simulate mutations under the GTR model with mean substitution rates of $5 \times 10^{-7}$, $5 \times 10^{-8}$, and $5 \times 10^{-9}$ per bp per generation. We vary mutation rates across sites and genomic regions in a 2-level approach. We divide the genomes into regions with a mean length of $10^4$ and multiply the mutation rates of each region by a random multiplier drawn from a Gamma distribution with a relatively high variance; within each region, we also multiply the mutation rate of *each position* from a Gamma distribution with a 20 times lower variance. The (inverse of the) variance itself is drawn from a log-normal distribution *per each replicate*.

Under the default condition, we let the mean population size be $10^5$ and the mean substitution rate be $5 \times 10^{-8}$; we sample a single copy per diploid genome (Table S2). To benchmark reconstruction methods under various conditions, we increase the population size to 10X and change the mutation rate to 0.1X and 10X. We also simulated unphased data from diploid genomes. For the scalability study, we simulate 2048 scaffold sequences of length $10^6$ bps (Table S3).

**Alternative methods**

On the simulated data, we ran alternative methods as follows. The exact versions, environment, and parallelization option used for each reconstruction tool are listed in Table S4. The exact commands are listed in the supplement.

By default, we ran RAxML-ng on the concatenated alignments under the GTR+G model with only one round using a parsimonious starting tree. For diploid genomes, we use the GT-GTR4+G model. We similarly ran SVDQuartets using all quartets of species. For diploid genomes, we sub-sample 10% of the quartets due to memory limitations.

We ran CASTER in specific ways on simulated data. To achieve a fair comparison with RAxML-ng, which was used with one starting tree, we ran CASTER with only one greedy tree and no tree merging step. Note that this was only for simulations; on real datasets, we used the default method of computing eight greedy trees and merging them, unless otherwise stated.

We ran wASTRAL differently from other methods. We (i) first partition the alignments into 2500-bp blocks; (ii) next, use RAxML-ng (default starting trees) to reconstruct a gene tree per block using only the first 500 bps; (iii) then, use IQ-Tree to annotate all gene trees with approximate Bayesian support values; (iv) finally, use wASTRAL-hybrid to infer species trees using annotated gene trees. The selection of 500 out of 2500 bp is to minimize recombinations within loci and increase independence across loci.

We ran BPP by first partitioning the alignments into 5000-bp blocks and then taking the first 200 bps per block as the input. The MCMC chain length is set to be 150,000 after a 100,000 burn-in. Due to the limited parallel performance of BPP, we only ran BPP on one CPU (8 physical cores) with a wall-time of 25 *days*. We ran snapper by first partitioning the alignments into 2500-bp blocks and then taking the first 50 bps per block as the input. The MCMC chain length is set to be 150,000 after a 100,000 burn-in.

#### Sliding-window analysis

##### Removing low-coverage regions

To create Fig. 3B-E, we assessed the CASTER scores for both main and alternative topologies across all branches. This was done over window sizes of 10k, 50k, 200k, 1M, and 5Mbps. Prior to this, we excluded low-coverage regions, as these areas are often associated with poor alignment quality. Note that these sites are included in the species tree analyses and only removed for those visualizations and statistical tests. The process for removing low-coverage regions is as follows:

1. Computing coverage: For each quadripartition $(P_1, P_2, P_3, P_4)$ corresponding to an internal branch of the species tree and every non-overlapping 10k window $\{i, i+1, \ldots i+$

9999}, we defined the coverage $C_i(P_1, P_2, P_3, P_4)$ as:

$$C_i(P_1, P_2, P_3, P_4) = \ln \sum_{j=i}^{i+9999} \sum_{(a,b,c,d)\in P_1\times P_2\times P_3\times P_4} \delta_a^j \delta_b^j \delta_c^j \delta_d^j,$$

where $\delta_a^j = 1$ if $p_a^j \in \{A, C, G, T\}$, and $\delta_a^j = 0$ otherwise. The same applies to $\delta_b^j, \delta_c^j$, and $\delta_d^j$. Thus, $\delta_a^j = 0$ denotes either a gap or an ambiguous letter. For each quadripartition $(P_1, P_2, P_3, P_4)$, we computed the set $C = \{C_i(P_1, P_2, P_3, P_4)\}$ across all non-overlapping 10k windows ($i \in \{0, 10000, \ldots\}$).

2. Filtering low-coverage 10kbp windows: We then removed 10kbp regions in $C$ that were at least 1.5 times the interquartile range (IQR) below the first quartile (Q1).

3. Excluding larger low-coverage windows: Finally, for each window size of 50k, 200k, 1M, and 5Mbp, we excluded the window if 3, 10, 50, or 250 of the respective 10kbp sub-windows within it were removed, depending on the size of the window under consideration.

**Normalized CASTER score**

To address the influence of gaps, ambiguous sites, and removed regions, we defined a normalized score for the topology $P_1P_2|P_3P_4$ on the window $[\alpha, \beta]$. This normalized score, accounting for all unaffected quartets, is determined using the following formula:

$$\bar{w}_{[\alpha,\beta]}(P_1, P_2; P_3, P_4) = \frac{\sum_{i=\alpha}^{\beta} \sum_{(a,b,c,d)\in P_1\times P_2\times P_3\times P_4} \delta_a^i \delta_b^i \delta_c^i \delta_d^i w_i(ab|cd)}{\sum_{i=\alpha}^{\beta} \sum_{(a,b,c,d)\in P_1\times P_2\times P_3\times P_4} \delta_a^i \delta_b^i \delta_c^i \delta_d^i}$$

In Fig. 3BE, for each branch corresponding to topology $P_1P_2|P_3P_4$, we displayed $\bar{w}_{[\alpha,\beta]}(P_1, P_2; P_3, P_4)$, $\bar{w}_{[\alpha,\beta]}(P_1, P_3; P_2, P_4)$, and $\bar{w}_{[\alpha,\beta]}(P_1, P_4; P_2, P_3)$ proportionally, ensuring a minimum floor value of zero.

##### Introgression signal detection

In the creation of Fig. 3CD, we computed p-values for the presence of incomplete lineage sorting (ILS)-like signals for each quadripartition with the topology $P_1P_2|P_3P_4$, considering various window sizes $[\alpha, \beta]$.

1. Computing total normalized score: We defined the total normalized score for the branch corresponding to $P_1P_2|P_3P_4$ within window $[\alpha, \beta]$ as the sum of all three topologies:
$$\begin{aligned}\bar{\sigma}_{[\alpha,\beta]}(P_1, P_2, P_3, P_4) =& \bar{w}_{[\alpha,\beta]}(P_1, P_2; P_3, P_4) \\ &+ \bar{w}_{[\alpha,\beta]}(P_1, P_3; P_2, P_4) + \bar{w}_{[\alpha,\beta]}(P_1, P_4; P_2, P_3)\end{aligned}$$

2. Small windows analysis (10k/50k/200kbps): For smaller windows, we *empirically* found that in scenarios with only ILS as true source of discordance, the total scores tend to follow an exponential distribution. In particular, close to extremely high scores, we observed:
$$\mathbb{P}\Big(\bar{\sigma}_{[\alpha,\beta]}(P_1, P_2, P_3, P_4) > \sigma\Big) \approx 0.05 \times 5^{-\frac{\sigma-\sigma_{95}}{\sigma_{99}-\sigma_{95}}}, \tag{7}$$
for $\sigma > \sigma_{95}$, where $\sigma_{95}$ and $\sigma_{99}$ denote the 95th and 99th percentile of all $\bar{\sigma}_{[\cdot,\cdot]}(P_1, P_2, P_3, P_4)$ for the same window size (Fig. S6). Therefore, the right-hand side of (7) gives an approximate p-value under the null hypothesis (i.e., ILS-only discordance); these were computed for the two alternative topologies. Windows with $p < 0.01$ were reported after Benjamini-Hochberg correction.

3. Large windows analysis (1Mbps): For larger windows, we computed the median ($\bar{\sigma}_{\text{MED}}$) and median absolute deviation ($\bar{\sigma}_{\text{MAD}}$) of $\bar{\sigma}_{[\cdot,\cdot]}(P_1, P_2, P_3, P_4)$. We calculated two p-values using the normal distribution for the two alternative topologies using
$$F_{\text{norm}}\Big(-\frac{0.6745\big(\bar{w}_{[\alpha,\beta]}(P_1, P_3; P_2, P_4) - \bar{\sigma}_{\text{MED}}\big)}{\bar{\sigma}_{\text{MAD}}}\Big) \text{ and}$$
$$F_{\text{norm}}\Big(-\frac{0.6745\big(\bar{w}_{[\alpha,\beta]}(P_1, P_4; P_2, P_3) - \bar{\sigma}_{\text{MED}}\big)}{\bar{\sigma}_{\text{MAD}}}\Big)$$

where $F_{\text{norm}}$ denotes the cumulative distribution function (CDF) of the standard normal distribution. The justification is that with a sufficiently large window size, it is reasonable to assume that sites spatially apart are nearly independent. Therefore, we can assume that $\bar{\sigma}$, the average of a particular site property over a large number of sites, approximately follows a normal distribution by the central limit theorem under weak dependence. We reported windows with $p < 0.01$ after Benjamini-Hochberg correction.

### Supplementary Text

#### Commands

CASTER (site and pair):

```
$PROGRAM_NAME -C -t $THREADS $MSA
```

wASTRAL (gene trees):

```
raxml-ng --msa $MSA --model GTR+G --threads 1
iqtree2 -s $MSA -te $MSA.raxml.bestTree \
    -m GTR+G4 -abayes -nt 1
```

wASTRAL (species tree):

```
astral-hybrid -B -C -t $THREADS $GENE_TREES
```

SVDQuartets (haploid, single individual):

```
svdquartets nthreads=$THREADS evalQuartets=all seed=5000
```

SVDQuartets (unphased diploid):

```
svdquartets nthreads=$THREADS evalQuartets=random \
    nquartets=105597360 taxpartition=species seed=5000
```

SVDQuartets (multiple individuals):

```
svdquartets nthreads=$THREADS evalQuartets=all \
    taxpartition=species seed=5000
```

RAxML-ng (haploid, single individual):

```
raxml-ng --tree pars{1} --msa $MSA --model GTR+G \
--threads $THREADS
```

RAxML-ng (unphased diploid):

```
raxml-ng --tree pars{1} --msa $MSA \
    --model GTGTR4+G --threads $THREADS
```

RAxML-ng (multiple individuals):

```
raxml-ng --tree pars{1} --msa $MSA --model GTR+G \
    --threads $THREADS --tree-constraint \
    $SPECIES_DELIMITATION
```

BPP (control file):

```
seed = 2333
seqfile = $MSA
Imapfile = $IMAP_FILE
outfile = $OUT_FILE
mcmcfile = $MCMC_FILE
speciesdelimitation = 0
speciestree = 1  0.4  0.2  0.1
speciesmodelprior = 1
species&tree = 21  T0 T10 T20 T30 T40 T50 T60 T70
    T80 T90 T100 T110 T120 T130 T140 T150 T160 T170
    T180 T190 T200
2 2 2 2 2 2 2 2 2 2 2 2 2 2 2 2 2 2 2 2 2
(((T60,((((T160,T200),T20),T180),((T140,(T90,T170)),
    (T80,T190)))),(((T120,T100),((T70,T30),T150)),
    ((T10,T50),(T130,(T110,T40))))),T0);
phase = 0 0 0 0 0 0 0 0 0 0 0 0 0 0 0 0 0 0 0 0 0
usedata = 1
nloci = 1000
model = gtr
cleandata = 0
thetaprior = 3 0.2
tauprior = 3 0.3
finetune =  1: 5 0.001 0.001  0.001 0.3 0.33 1.0
print = 1 0 0 0
burnin = 100000
sampfreq = 5
nsample = 150000
Threads = 8
```

snapper (control file):

```
<beast ... required="BEAST.base■v2.7.3:snapper
■■■■v1.1.0:SNAPP■v1.6.1" version="2.7">
...
<run id="mcmc" spec="MCMC" chainLength="150000"
  preBurnin="100000" storeEvery="1000">
<state id="state" spec="State" storeEvery="5000">
...
```

Placental mammal whole genome alignment:

```
caster-site -C -t 64 -o chr1.nw chr1.fa
caster-site -r 0 -s 0 -t 128 -f list -g chr1.nw \
    -o whole_genome.nw chr_list.txt
```

#### Supplementary equations

Equivalent of Equation (6) for the F84 model:

$$
\begin{aligned}
&W_i(P_1; P_2, P_3) \\
&= \left( 2\pi_A\pi_G \left( \binom{\mathbf{\Sigma}_1[i,A]}{2} + \binom{\mathbf{\Sigma}_1[i,G]}{2} \right) - (\pi_A^2 + \pi_G^2)\mathbf{\Sigma}_1[i,A]\mathbf{\Sigma}_1[i,G] \right) \\
&\qquad \Big( 2\pi_C\pi_T \big( \mathbf{\Sigma}_2[i,C]\mathbf{\Sigma}_3[i,C] + \mathbf{\Sigma}_2[i,T]\mathbf{\Sigma}_3[i,T] \big) \\
&\qquad\qquad - (\pi_C^2 + \pi_T^2)\big( \mathbf{\Sigma}_2[i,C]\mathbf{\Sigma}_3[i,T] + \mathbf{\Sigma}_2[i,T]\mathbf{\Sigma}_3[i,C] \big) \Big) \\
&+ \left( 2\pi_C\pi_T \left( \binom{\mathbf{\Sigma}_1[i,C]}{2} + \binom{\mathbf{\Sigma}_1[i,T]}{2} \right) - (\pi_C^2 + \pi_T^2)\mathbf{\Sigma}_1[i,C]\mathbf{\Sigma}_1[i,T] \right) \\
&\qquad \Big( 2\pi_A\pi_G \big( \mathbf{\Sigma}_2[i,A]\mathbf{\Sigma}_3[i,A] + \mathbf{\Sigma}_2[i,G]\mathbf{\Sigma}_3[i,G] \big) \\
&\qquad\qquad - (\pi_A^2 + \pi_G^2)\big( \mathbf{\Sigma}_2[i,A]\mathbf{\Sigma}_3[i,G] + \mathbf{\Sigma}_2[i,G]\mathbf{\Sigma}_3[i,A] \big) \Big)
\end{aligned}
\tag{8}
$$

Equivalent of Equation (6) for the GTR model:

Notice that under the GTR model, the entries of the matrices are $\mathbf{\Sigma}_j[i,AA]$, $\mathbf{\Sigma}_j[i,AC]$, $\mathbf{\Sigma}_j[i,AG]$, ..., $\mathbf{\Sigma}_j[i,TT]$. To simplify notations, we denote

$$RN_j^i = \sum_{X\in\{A,G\}} \sum_{Z\in\{A,G,C,T\}} \mathbf{\Sigma}_j[i,XZ] \ ,$$

ditto for $NR_j^i, YN_j^i, \ldots NN_j^i$.

$$\begin{aligned}
&W_i(P_1;P_2,P_3) = \\
&\sum_{(X,Z)\in\{(R,Y),(M,K),(W,S)\}} (XN_1^i)(ZN_1^i)\big((NX_2^i)(NZ_3^i)+(NZ_2^i)(NX_3^i)\big) \\
&\qquad +(NX_1^i)(NZ_1^i)\big((XN_2^i)(ZN_3^i)+(ZN_2^i)(XN_3^i)\big) \\
&\qquad -4\pi_X\pi_Z(XN_1^i)(ZN_1^i)(NN_2^i)(NN_3^i) \\
&\qquad -4\pi_X\pi_Z\binom{NN_1^i}{2}\big((XN_2^i)(ZN_3^i)+(ZN_2^i)(XN_3^i)\big)
\end{aligned} \tag{9}$$

#### Proofs

**Notation:** Across proofs, we use these additional notations. For each tree topology $S_*$, we let $\mathbf{L}(S_*)$ denote the set of leaf labels of $S_*$. We let $\mathcal{Q}(S_*) = \{\mathbf{q} : \mathbf{q} \subseteq \mathbf{L}(S_*), |\mathbf{q}| = 4\}$ denote the set of quartets of leaves. For each quartet $\mathbf{q}$, we let $S_* \upharpoonright \mathbf{q}$ denote the quartet tree topology of $S_*$ restricted to $\mathbf{q}$. Notice that $\mathbf{T}(S_*) = \{S_* \upharpoonright \mathbf{q} : \mathbf{q} \in \mathcal{Q}(S_*)\}$.

For the true species tree $\mathbf{S}$ and any other topology $S_*$, let $\Delta\mathcal{Q} = \{\mathbf{q} : \mathbf{S} \upharpoonright \mathbf{q} \neq S_* \upharpoonright \mathbf{q}, \mathbf{q} \in \mathcal{Q}(S_*)\}$ denote the set of quartets for which $\mathbf{S}$ and $S_*$ differ in topology.

*Proof of Theorem 1.* To prove that maximizing $W(S_*) = \sum_{T \in \mathbf{T}(S_*)} \sum_{i=1}^{k} w_i(T)$ consistently estimates the true species tree topology $\mathbf{S}$, we need to show the following premises:

1. For all $T_1, T_2$ and $i \neq j$, $w_i(T_1)$ and $w_j(T_2)$ are independent.
2. For all $T \in \mathbf{T}(\mathbf{S})$ and $i$, $|w_i(T)|$ is bounded.
3. For all $T \in \mathbf{T}(\mathbf{S})$ and $i$, $\mathbb{E}\left[w_i(ab|cd)\right] > \mathbb{E}\left[w_i(ac|bd)\right] = \mathbb{E}\left[w_i(ad|bc)\right]$, where $T = ab|cd$.

The first premise follows from the independence assumption stated in Models and notations, and the second premise can be easily verified from Figs. 1cd and S1. The third premise is formally stated in Proposition 1 and will be proved after this proof.

Recalling the notations shown above, the difference in objectives is:

$$\begin{aligned} W(\mathbf{S}) - W(S_*) &= \sum_{i=1}^{k} \sum_{\mathbf{q} \in \mathcal{Q}(\mathbf{S})} w_i(\mathbf{S} \upharpoonright \mathbf{q}) - w_i(S_* \upharpoonright \mathbf{q}) \\ &= \sum_{i=1}^{k} \sum_{\mathbf{q} \in \Delta\mathcal{Q}} w_i(\mathbf{S} \upharpoonright \mathbf{q}) - w_i(S_* \upharpoonright \mathbf{q}) \end{aligned} \tag{10}$$

From the second premise, it follows that $\left|\sum_{\mathbf{q} \in \Delta\mathcal{Q}} w_i(\mathbf{S} \upharpoonright \mathbf{q}) - w_i(S_* \upharpoonright \mathbf{q})\right|$ is bounded. From the third premise, we show that

$$\mathbb{E}\Big[\sum_{\mathbf{q} \in \Delta\mathcal{Q}} w_i(\mathbf{S} \upharpoonright \mathbf{q}) - w_i(S_* \upharpoonright \mathbf{q})\Big] > 0.$$

With the first premise, we can conclude that as $k \to +\infty$,

$$\mathbb{P}\big[W(\mathbf{S}) - W(S_*) > 0\big] = 1 \tag{11}$$

by the law of large numbers. □

**Proposition 1.** *For the true species tree* S *of four leaves with topology* $ab|cd$*,*

$$\forall i, \mathbb{E}\big[w_i(ab|cd)\big] > \mathbb{E}\big[w_i(ac|bd)\big] = \mathbb{E}\big[w_i(ac|bd)\big] \ . \tag{12}$$

*Proof.* We start with a premise, which will be formally stated and proved in Proposition 2:

Conditional on a given gene tree $\mathbf{G}_i$ with topology $ab|cd$,

$$\mathbb{E}\big[w_i(ab|cd)\big|\mathbf{G}_i\big] - \alpha\big(L_T(\mathbf{G}_i)\big)\beta\big(L_I(\mathbf{G}_i)\big) = \mathbb{E}\big[w_i(ac|bd)\big|\mathbf{G}_i\big] = \mathbb{E}\big[w_i(ad|bc)\big|\mathbf{G}_i\big] \ , \tag{13}$$

where $\alpha : \mathbb{R}_+ \to \mathbb{R}_+$ and $\beta : \mathbb{R}_+ \to \mathbb{R}_+$ are two functions independent of $\mathbf{G}_i$; $L_T(\mathbf{G}_i)$ denotes the total terminal branch length of $\mathbf{G}_i$ and $L_I(\mathbf{G}_i)$ denotes the internal branch length of $\mathbf{G}_i$. In other words, the difference between expected scores of alternative topologies and the main topology *only* depends on the total terminal branch length (in substitution units) and the internal branch length.

It is sufficient to prove the following equation is greater than zero:

$$\mathbb{E}\big[w_i(ab|cd) - w_i(ac|bd)\big] = \int \mathbb{E}\big[w_i(ab|cd) - w_i(ac|bd)\big|\mathbf{G}_i\big] f(\mathbf{G}_i) d\mathbf{G}_i \tag{14}$$

where $\mathbf{G}_i$ is integrated over all possibilities, and $f(\cdot)$ is the probability density function of $\mathbf{G}_i$.

We divide this problem into two cases by the rooted topology of $\mathbf{S}$.

Case 1, S has an unbalanced topology:

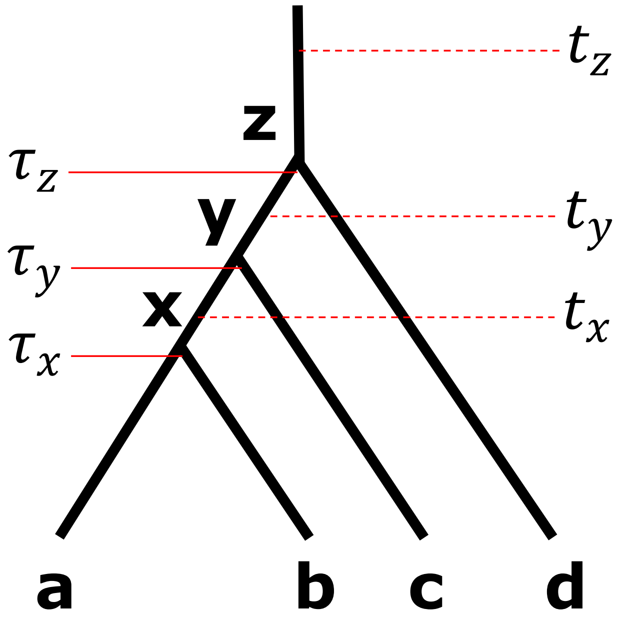


Without loss of generality, we assume $a$ and $b$ are sisters, and $d$ is the outgroup. Let $x$, $y$, and $z$ be the most recent common ancestor of $(a, b)$, $(a, c)$, and $(a, d)$, respectively. Recall that $\tau_x$, $\tau_y$, and $\tau_z$ denote the ages of $x$, $y$, and $z$, respectively.

We define the following branch lengths in substitution units:

- $l_a = \int_0^{\tau_x} \mu^i_{(a,x)}(t)dt$, the branch length of $(a, x)$;
- $l_b = \int_0^{\tau_x} \mu^i_{(b,x)}(t)dt$, the branch length of $(b, x)$;
- $l_c = \int_0^{\tau_y} \mu^i_{(c,y)}(t)dt$, the branch length of $(c, y)$;
- $l_d = \int_0^{\tau_z} \mu^i_{(d,z)}(t)dt$, the branch length of $(d, z)$.

We also define:

- $l_x(t_x) = \int_{\tau_x}^{t_x} \mu^i_{(x,y)}(t)dt$ for any $\tau_x < t_x < \tau_y$;
- $l_y(t_y) = \int_{\tau_y}^{t_y} \mu^i_{(y,z)}(t)dt$ for any $\tau_y < t_y < \tau_z$;
- $l_z(t_z) = \int_{\tau_z}^{t_z} \mu^i_{(z,\emptyset)}(t)dt$ for any $\tau_z < t_z$.

Notice that we can divide all possible $\mathbf{G}_i$'s into two categories:

1. Sets of $\mathbf{G}_i$'s with the same coalescent times and hence $L_T(\mathbf{G}_i)$, $L_I(\mathbf{G}_i)$, and $f(\mathbf{G}_i)$, but different unrooted topologies (cases a–e shown in the Figure below). In this case, for each set $\mathbf{G}'$, due to Proposition 2 and symmetry,

$$\sum_{\mathbf{G}_i \in \mathbf{G}'} \mathbb{E}\big[w_i(ab|cd) - w_i(ac|bd)\big|\mathbf{G}_i\big] f(\mathbf{G}_i) = 0$$

2. The rest of $\mathbf{G}_i$'s where $a$ and $b$ avoid deep coalescence (case f in the figure below) and thus all have the same topology $ab|cd$; here, by Proposition 2,

$$\mathbb{E}\big[w_i(ab|cd) - w_i(ac|bd)\big|\mathbf{G}_i\big] f(\mathbf{G}_i) > 0.$$

Below are the illustrations of both categories and their perspective

$$\mathbb{E}\big[w_i(ab|cd) - w_i(ac|bd)\big|\mathbf{G}_i\big] \text{ values}$$

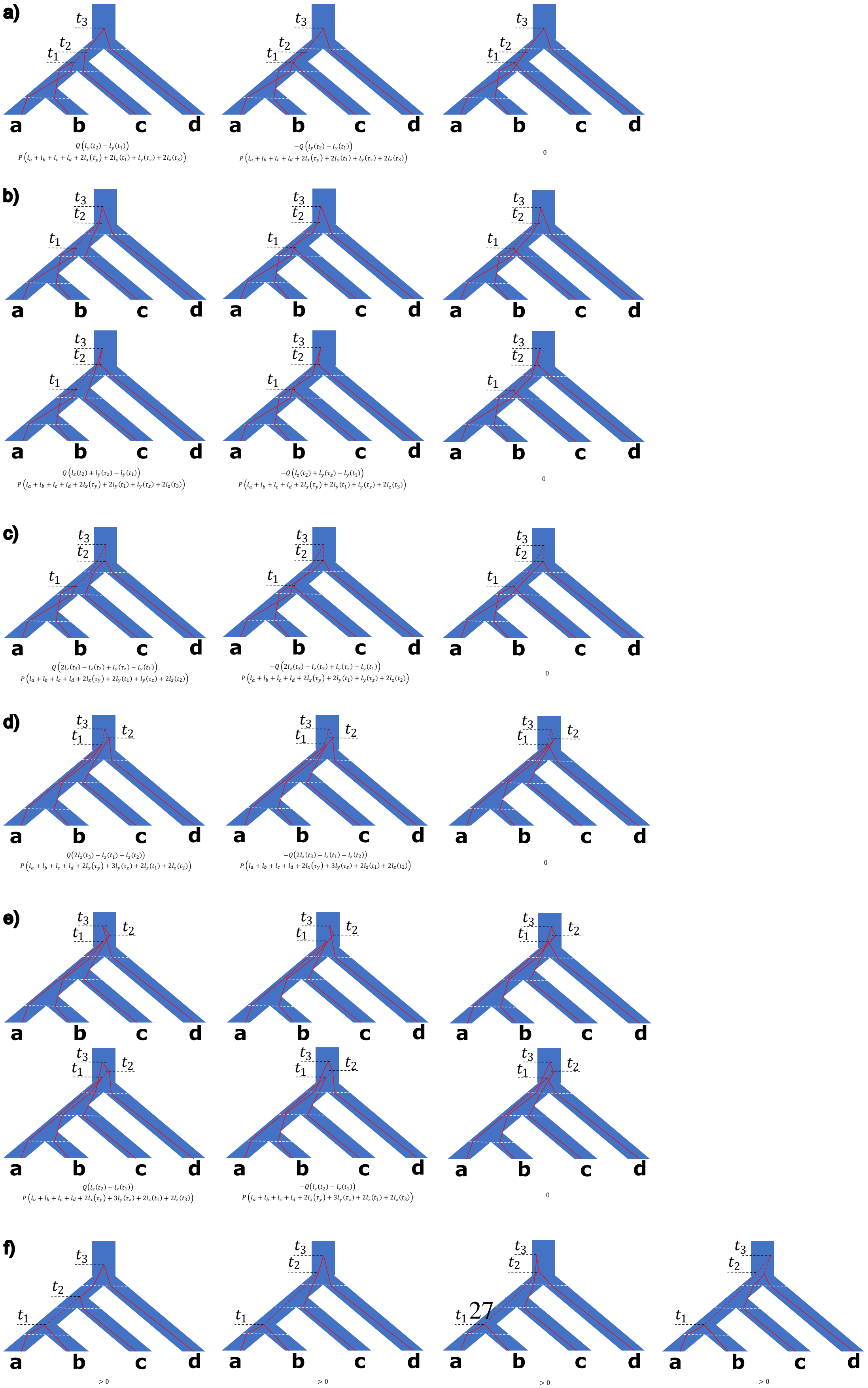
a)
$t_3$ $t_2$ $t_1$
a b c d
$Q\left(l_y(t_2) - l_y(t_1)\right)$
$P\left(l_a + l_b + l_c + l_d + 2l_x(\tau_y) + 2l_y(t_1) + l_y(\tau_z) + 2l_z(t_3)\right)$
$-Q\left(l_y(t_2) - l_y(t_1)\right)$
$P\left(l_a + l_b + l_c + l_d + 2l_x(\tau_y) + 2l_y(t_1) + l_y(\tau_z) + 2l_z(t_3)\right)$
0
b)
$Q\left(l_z(t_2) + l_y(\tau_z) - l_y(t_1)\right)$
$P\left(l_a + l_b + l_c + l_d + 2l_x(\tau_y) + 2l_y(t_1) + l_y(\tau_z) + 2l_z(t_3)\right)$
$-Q\left(l_z(t_2) + l_y(\tau_z) - l_y(t_1)\right)$
$P\left(l_a + l_b + l_c + l_d + 2l_x(\tau_y) + 2l_y(t_1) + l_y(\tau_z) + 2l_z(t_3)\right)$
0
c)
$Q\left(2l_z(t_3) - l_z(t_2) + l_y(\tau_z) - l_y(t_1)\right)$
$P\left(l_a + l_b + l_c + l_d + 2l_x(\tau_y) + 2l_y(t_1) + l_y(\tau_z) + 2l_z(t_2)\right)$
$-Q\left(2l_z(t_3) - l_z(t_2) + l_y(\tau_z) - l_y(t_1)\right)$
$P\left(l_a + l_b + l_c + l_d + 2l_x(\tau_y) + 2l_y(t_1) + l_y(\tau_z) + 2l_z(t_2)\right)$
0
d)
$Q(2l_z(t_3) - l_z(t_1) - l_z(t_2))$
$P\left(l_a + l_b + l_c + l_d + 2l_x(\tau_y) + 3l_y(\tau_z) + 2l_z(t_1) + 2l_z(t_2)\right)$
$-Q(2l_z(t_3) - l_z(t_1) - l_z(t_2))$
$P\left(l_a + l_b + l_c + l_d + 2l_x(\tau_y) + 3l_y(\tau_z) + 2l_z(t_1) + 2l_z(t_2)\right)$
0
e)
$Q(l_z(t_2) - l_z(t_1))$
$P\left(l_a + l_b + l_c + l_d + 2l_x(\tau_y) + 3l_y(\tau_z) + 2l_z(t_1) + 2l_z(t_3)\right)$
$-Q(l_z(t_2) - l_z(t_1))$
$P\left(l_a + l_b + l_c + l_d + 2l_x(\tau_y) + 3l_y(\tau_z) + 2l_z(t_1) + 2l_z(t_3)\right)$
0
f)
> 0
> 0
> 0
> 0

Thus, in total:

$$\mathbb{E}\big[w_i(ab|cd) - w_i(ac|bd)\big] = \int \mathbb{E}\big[w_i(ab|cd) - w_i(ac|bd)\big|\mathbf{G}_i\big] f(\mathbf{G}_i) d\mathbf{G}_i > 0. \qquad (15)$$

Case 2, S has a balanced topology:

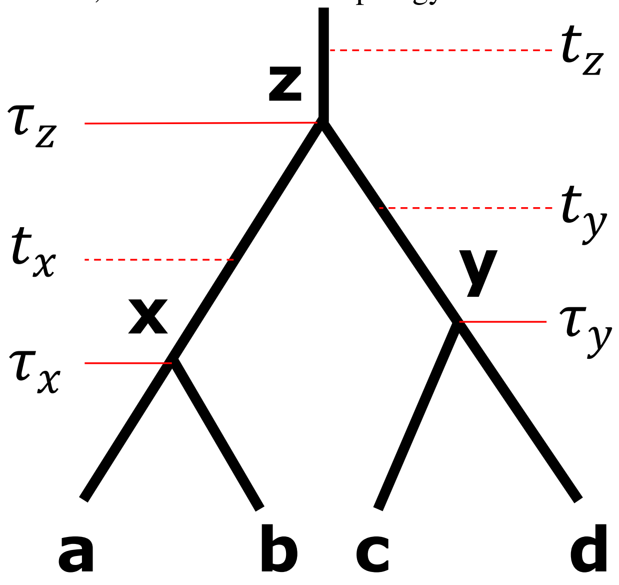


Without loss of generality, we assume $a$ and $b$ are sisters, and $c$ and $d$ are sisters. Let $x$, $y$, and $z$ be the most recent common ancestor of $(a, b)$, $(c, d)$, and $(a, d)$, respectively. Recall that $\tau_x$, $\tau_y$, and $\tau_z$ denote the ages of $x$, $y$, and $z$, respectively.

We define the following branch lengths in substitution units:

- $l_a = \int_0^{\tau_x} \mu^i_{(a,x)}(t)dt$, the branch length of $(a, x)$;
- $l_b = \int_0^{\tau_x} \mu^i_{(b,x)}(t)dt$, the branch length of $(b, x)$;
- $l_c = \int_0^{\tau_y} \mu^i_{(c,y)}(t)dt$, the branch length of $(c, y)$;
- $l_d = \int_0^{\tau_z} \mu^i_{(d,y)}(t)dt$, the branch length of $(d, y)$.

We also define:

- $l_x(t_x) = \int_{\tau_x}^{t_x} \mu^i_{(x,z)}(t)dt$ for any $\tau_x < t_x < \tau_z$;

- $l_y(t_y) = \int_{\tau_y}^{t_y} \mu^i_{(y,z)}(t)dt$ for any $\tau_y < t_y < \tau_z$;
- $l_z(t_z) = \int_{\tau_z}^{t_z} \mu^i_{(z,\emptyset)}(t)dt$ for any $\tau_z < t_z$.

Notice that we can also divide all possible $\mathbf{G}_i$'s into two categories:

1. Sets of $\mathbf{G}_i$'s with the same $L_T(\mathbf{G}_i)$, $L_I(\mathbf{G}_i)$, and $f(\mathbf{G}_i)$, but different unrooted quartet topologies (cases a and b of the figure below). In this case, for each set $\mathbf{G}'$,

$$\sum_{\mathbf{G}_i \in \mathbf{G}'} \mathbb{E}\big[w_i(ab|cd) - w_i(ac|bd)\big|\mathbf{G}_i\big] f(\mathbf{G}_i) = 0.$$

2. The rest of $\mathbf{G}_i$'s all have the same topology $ab|cd$ (e.g. as shown in panel c below), and by Proposition 2,

$$\mathbb{E}\big[w_i(ab|cd) - w_i(ac|bd)\big|\mathbf{G}_i\big] f(\mathbf{G}_i) > 0.$$

Below are the illustrations of both categories and their respective

$$\mathbb{E}\big[w_i(ab|cd) - w_i(ac|bd)\big|\mathbf{G}_i\big].$$

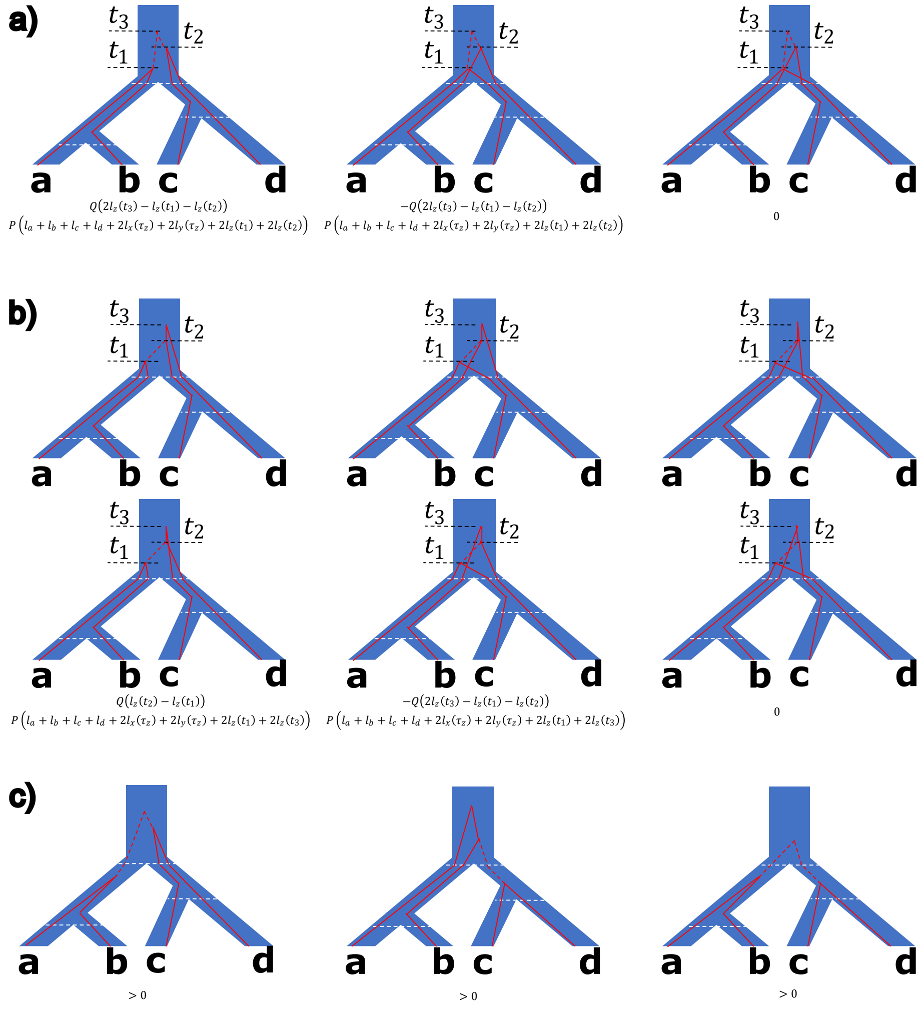


Thus, in total:

$$\mathbb{E}\big[w_i(ab|cd) - w_i(ac|bd)\big] = \int \mathbb{E}\big[w_i(ab|cd) - w_i(ac|bd)\big|\mathbf{G}_i\big] f(\mathbf{G}_i) d\mathbf{G}_i > 0. \tag{16}$$

$\square$

**Proposition 2.** *For the true gene tree $\mathbf{G}_i$ of four leaves with topology $ab|cd$, let $l_a$, $l_b$, $l_c$, $l_d$ denote the terminal branch lengths, and $l_x$ denote the internal branch length in substitution units, then*

$$\mathbb{E}\big[w_i(ab|cd)\big|\mathbf{G}_i\big] - \gamma e^{-\alpha(l_a+l_b+l_c+l_d)}\big(1 - e^{-\beta l_x}\big) = \mathbb{E}\big[w_i(ac|bd)\big|\mathbf{G}_i\big] \tag{17}$$

*for some $\alpha > 0$, $\beta > 0$, and $\gamma > 0$.*

*Proof.* We start with the simplest JC69 model:

$$Q = \begin{bmatrix} -1 & \frac{1}{3} & \frac{1}{3} & \frac{1}{3} \\ \frac{1}{3} & -1 & \frac{1}{3} & \frac{1}{3} \\ \frac{1}{3} & \frac{1}{3} & -1 & \frac{1}{3} \\ \frac{1}{3} & \frac{1}{3} & \frac{1}{3} & -1 \end{bmatrix}, e^{Qt} = \begin{bmatrix} \frac{1}{4}+\frac{3}{4}e^{-\frac{4}{3}t} & \frac{1}{4}-\frac{1}{4}e^{-\frac{4}{3}t} & \frac{1}{4}-\frac{1}{4}e^{-\frac{4}{3}t} & \frac{1}{4}-\frac{1}{4}e^{-\frac{4}{3}t} \\ \frac{1}{4}-\frac{1}{4}e^{-\frac{4}{3}t} & \frac{1}{4}+\frac{3}{4}e^{-\frac{4}{3}t} & \frac{1}{4}-\frac{1}{4}e^{-\frac{4}{3}t} & \frac{1}{4}-\frac{1}{4}e^{-\frac{4}{3}t} \\ \frac{1}{4}-\frac{1}{4}e^{-\frac{4}{3}t} & \frac{1}{4}-\frac{1}{4}e^{-\frac{4}{3}t} & \frac{1}{4}+\frac{3}{4}e^{-\frac{4}{3}t} & \frac{1}{4}-\frac{1}{4}e^{-\frac{4}{3}t} \\ \frac{1}{4}-\frac{1}{4}e^{-\frac{4}{3}t} & \frac{1}{4}-\frac{1}{4}e^{-\frac{4}{3}t} & \frac{1}{4}-\frac{1}{4}e^{-\frac{4}{3}t} & \frac{1}{4}+\frac{3}{4}e^{-\frac{4}{3}t} \end{bmatrix} \tag{18}$$

Let

$$\mathbf{a} = \begin{bmatrix} 1 \\ 0 \\ 0 \\ 0 \end{bmatrix}, \mathbf{g} = \begin{bmatrix} 0 \\ 1 \\ 0 \\ 0 \end{bmatrix}, \mathbf{c} = \begin{bmatrix} 0 \\ 0 \\ 1 \\ 0 \end{bmatrix}, \mathbf{t} = \begin{bmatrix} 0 \\ 0 \\ 0 \\ 1 \end{bmatrix}, \mathbf{\Pi} = \begin{bmatrix} 0.25 \\ 0.25 \\ 0.25 \\ 0.25 \end{bmatrix}, \tag{19}$$

and $P(\mathbf{v_a}, \mathbf{v_b}, \mathbf{v_c}, \mathbf{v_d})$ denote the probability that $p_a^i = \mathbf{v_a}, p_b^i = \mathbf{v_b}, p_c^i = \mathbf{v_c}, p_d^i = \mathbf{v_d}$ given $\mathbf{G}_i$, for all $\mathbf{v_a}, \mathbf{v_b}, \mathbf{v_c}, \mathbf{v_d} \in \{\mathbf{a}, \mathbf{g}, \mathbf{c}, \mathbf{t}\}$. In fact, $P(\mathbf{v_a}, \mathbf{v_b}, \mathbf{v_c}, \mathbf{v_d})$ can be computed using the following equation:

$$P(\mathbf{v_a}, \mathbf{v_b}, \mathbf{v_c}, \mathbf{v_d}) = \mathbf{\Pi}^\intercal \Big( e^{l_a Q}\mathbf{v_a} \odot e^{l_b Q}\mathbf{v_b} \odot \big( e^{l_x Q}(e^{l_c Q}\mathbf{v_c} \odot e^{l_d Q}\mathbf{v_d})\big)\Big) \tag{20}$$

According to Fig. 1C,

$$\begin{aligned} \mathbb{E}\big[w_i(ab|cd)\big|\mathbf{G}_i\big] &= 2 \sum_{\{\mathbf{x},\mathbf{y}\}\subseteq\mathbf{\Sigma}} P(\mathbf{x},\mathbf{x},\mathbf{y},\mathbf{y}) \\ &\quad - \sum_{\mathbf{x}\in\mathbf{\Sigma}} \sum_{\mathbf{y}\in\mathbf{\Sigma}\setminus\{\mathbf{x}\}} \sum_{\mathbf{z}\in\mathbf{\Sigma}\setminus\{\mathbf{x},\mathbf{y}\}} P(\mathbf{x},\mathbf{x},\mathbf{y},\mathbf{z}) + P(\mathbf{y},\mathbf{z},\mathbf{x},\mathbf{x}) \ . \end{aligned} \tag{21}$$

$$\begin{aligned} \mathbb{E}\big[w_i(ac|bd)\big|\mathbf{G}_i\big] &= 2 \sum_{\{\mathbf{x},\mathbf{y}\}\subseteq\mathbf{\Sigma}} P(\mathbf{x},\mathbf{y},\mathbf{x},\mathbf{y}) \\ &\quad - \sum_{\mathbf{x}\in\mathbf{\Sigma}} \sum_{\mathbf{y}\in\mathbf{\Sigma}\setminus\{\mathbf{x}\}} \sum_{\mathbf{z}\in\mathbf{\Sigma}\setminus\{\mathbf{x},\mathbf{y}\}} P(\mathbf{x},\mathbf{y},\mathbf{x},\mathbf{z}) + P(\mathbf{y},\mathbf{x},\mathbf{z},\mathbf{x}) \ , \end{aligned} \tag{22}$$

where $\mathbf{\Sigma} = \{\mathbf{a}, \mathbf{g}, \mathbf{c}, \mathbf{t}\}$. Since for any distinct $\mathbf{x}, \mathbf{y}, \mathbf{z}$, by symmetry,

$$P(\mathbf{x},\mathbf{x},\mathbf{y},\mathbf{y}) = P(\mathbf{a},\mathbf{a},\mathbf{c},\mathbf{c}) = P(\mathbf{c},\mathbf{c},\mathbf{a},\mathbf{a}) \text{ and } P(\mathbf{x},\mathbf{x},\mathbf{y},\mathbf{z}) = P(\mathbf{a},\mathbf{a},\mathbf{c},\mathbf{g}) \ , \tag{23}$$

it is sufficient to prove (17) correct under the JC69 model with the following patterns and weights:

| | | | | | | |
|---|---|---|---|---|---|---|
| **a** | A | C | A | A | C | G |
| **b** | A | C | A | A | G | C |
| **c** | C | A | C | G | A | A |
| **d** | C | A | G | C | A | A |
| **W** | +1 | | -1 | | | |

In this case,

$$\mathbb{E}\big[w_i(ab|cd)\big|\mathbf{G}_i\big] = P(\mathbf{a},\mathbf{a},\mathbf{c},\mathbf{c}) + P(\mathbf{c},\mathbf{c},\mathbf{a},\mathbf{a}) \\ - P(\mathbf{a},\mathbf{a},\mathbf{c},\mathbf{g}) - P(\mathbf{a},\mathbf{a},\mathbf{g},\mathbf{c}) - P(\mathbf{c},\mathbf{g},\mathbf{a},\mathbf{a}) - P(\mathbf{g},\mathbf{c},\mathbf{a},\mathbf{a}) \tag{24}$$

and

$$\mathbb{E}\big[w_i(ac|bd)\big|\mathbf{G}_i\big] = P(\mathbf{a},\mathbf{c},\mathbf{a},\mathbf{c}) + P(\mathbf{c},\mathbf{a},\mathbf{c},\mathbf{a}) \\ - P(\mathbf{a},\mathbf{c},\mathbf{a},\mathbf{g}) - P(\mathbf{a},\mathbf{g},\mathbf{a},\mathbf{c}) - P(\mathbf{c},\mathbf{a},\mathbf{g},\mathbf{a}) - P(\mathbf{g},\mathbf{a},\mathbf{c},\mathbf{a}) \ . \tag{25}$$

It can be mechanically verified that (Mathematica script provided)

$$\mathbb{E}\big[w_i(ab|cd) - w_i(ac|bd)\big|\mathbf{G}_i\big] = \frac{1}{8}e^{-\frac{4}{3}(l_a+l_b+l_c+l_d)}(1 - e^{-\frac{4}{3}l_x}) \ . \tag{26}$$

We omit the proof for F84 model, as it follows the same proof. For F84 model,

$$Q = \lambda \begin{bmatrix} * & \kappa\frac{\pi_G}{\pi_R} + \pi_G & \pi_C & \pi_T \\ \kappa\frac{\pi_A}{\pi_R} + \pi_A & * & \pi_C & \pi_T \\ \pi_A & \pi_G & * & \kappa\frac{\pi_T}{\pi_Y} + \pi_T \\ \pi_A & \pi_G & \kappa\frac{\pi_C}{\pi_Y} + \pi_C & * \end{bmatrix} \tag{27}$$

where $*$ are omitted for conciseness, and

$$\lambda = \frac{\pi_R \pi_Y}{(1 - \pi_A^2 - \pi_G^2 - \pi_C^2 - \pi_T^2)\pi_R\pi_Y + 2\kappa(\pi_A\pi_G\pi_Y + \pi_C\pi_T\pi_R)}. \tag{28}$$

It is also sufficient to prove (17) correct under the F84 model with the following patterns and weights:

| | | | | | | | | |
|---|---|---|---|---|---|---|---|---|
| **a** | R | Y | A | Y | R | C | R | Y |
| **b** | R | Y | A | Y | R | C | R | Y |
| **c** | Y | R | Y | A | C | R | Y | R |
| **d** | Y | R | Y | A | C | R | Y | R |
| **W** | $+\pi_A^2\pi_C^2$ | | $-\pi_R^2\pi_C^2$ | | $-\pi_A^2\pi_Y^2$ | | $+\pi_R^2\pi_Y^2$ | |

In this case,

$$\begin{aligned} &\mathbb{E}\left[w_i(ab|cd) - w_i(ac|bd)\middle|\mathbf{G}_i\right] \\ &\quad = 2\pi_A\pi_C\pi_G\pi_T\pi_R\pi_Y e^{-\lambda(1+\kappa)(l_a+l_b+l_c+l_d)}\left(1 - e^{-\lambda l_x}\right). \end{aligned} \tag{29}$$

For GTR model, again by symmetry, it is sufficient to prove (17) correct under the GTR model with the following patterns and weights:

| | | | | | | | | | | | | |
|---|---|---|---|---|---|---|---|---|---|---|---|---|
| **a** | RN | RN | YN | YN | NR | NR | NY | NY | RN | YN | NN | NN |
| **b** | YN | YN | RN | RN | NY | NY | NR | NR | YN | RN | NN | NN |
| **c** | NR | NY | NR | NY | RN | YN | RN | YN | NN | NN | RN | YN |
| **d** | NY | NR | NY | NR | YN | RN | YN | RN | NN | NN | YN | RN |
| **W** | $+1$ | | | | | | | | $-4\pi_R\pi_Y$ | | | |

We have

$$Q = \frac{1}{4\pi_R\pi_Y}\begin{bmatrix} -2\pi_Y & \pi_Y & \pi_Y & 0 \\ \pi_R & -1 & 0 & \pi_Y \\ \pi_R & 0 & -1 & \pi_Y \\ 0 & \pi_R & \pi_R & -2\pi_R \end{bmatrix} \tag{30}$$

for states $RR$, $RY$, $YR$, and $YY$.

We can still follow the same proof to get

$$\begin{aligned} &\mathbb{E}\big[w_i(ab|cd) - w_i(ac|bd)\big|\mathbf{G}_i\big] \\ &= 8\pi_R^2\pi_Y^2 e^{-\frac{1}{4\pi_R\pi_Y}(l_a+l_b+l_c+l_d)}(1 - e^{-\frac{1}{2\pi_R\pi_Y}l_x})\ . \end{aligned} \tag{31}$$

Here, we provide an alternative proof instead:

Let $\mathbb{P}(p_\mathbf{a} = R)$ denote the probability that the species $a$ has letter $R$ (A or G) at a site under $\mathbf{G}_i$; and let $\mathbb{P}(p_\mathbf{a} = RN)$ denote the probability that the pair of sites for species $a$ are $RN$ ($\{A, G\} \times \{A, G, C, T\}$).

For GTR model of only two states (R and Y), the (unnormalized) substitution matrix is very simple:

$$Q = \begin{bmatrix} -\pi_Y & \pi_Y \\ \pi_R & \pi_R \end{bmatrix} \tag{32}$$

Thus, we can easily compute the joint probabilities

$$\mathbb{P}(p_\mathbf{a} = R, p_\mathbf{b} = Y) = \pi_R\pi_Y(1 - e^{-\lambda(l_a+l_b)}) \tag{33}$$

and

$$\mathbb{P}(p_\mathbf{a} = R, p_\mathbf{c} = Y) = \pi_R\pi_Y(1 - e^{-\lambda(l_a+l_x+l_c)}) \tag{34}$$

for some $\lambda > 0$. And trivially,

$$\mathbb{P}(p_{\mathbf{a}} = RN, p_{\mathbf{b}} = YN) = \pi_R \pi_Y (1 - e^{-\lambda(l_a + l_b)}) \ . \tag{35}$$

and thus

$$\mathbb{P}(p_{\mathbf{a}} = RN, p_{\mathbf{b}} = YN, p_{\mathbf{c}} = NN, p_{\mathbf{d}} = NN) = \pi_R \pi_Y (1 - e^{-\lambda(l_a + l_b)}) \ . \tag{36}$$

Since

$$\mathbb{P}(p_{\mathbf{a}} = RN, p_{\mathbf{b}} = YN) = \pi_R \pi_Y (1 - e^{-\lambda(l_a + l_b)}), \tag{37}$$

$$\mathbb{P}(p_{\mathbf{c}} = NR, p_{\mathbf{d}} = NY) = \pi_R \pi_Y (1 - e^{-\lambda(l_c + l_d)}) \ , \tag{38}$$

and the first and the second site are independent, we have

$$\mathbb{P}(p_{\mathbf{a}} = RN, p_{\mathbf{b}} = YN, p_{\mathbf{c}} = NR, p_{\mathbf{d}} = NY) = \pi_R^2 \pi_Y^2 (1 - e^{-\lambda(l_a + l_b)})(1 - e^{-\lambda(l_c + l_d)}) \ . \tag{39}$$

According to the weights above,

$$\begin{aligned}
\mathbb{E}\big[w_i(ab|cd)\big|\mathbf{G}_i\big] &= \mathbb{P}(p_{\mathbf{a}} = RN, p_{\mathbf{b}} = YN, p_{\mathbf{c}} = NR, p_{\mathbf{d}} = NY) \\
&+ \mathbb{P}(p_{\mathbf{a}} = RN, p_{\mathbf{b}} = YN, p_{\mathbf{c}} = NY, p_{\mathbf{d}} = NR) \\
&+ \mathbb{P}(p_{\mathbf{a}} = YN, p_{\mathbf{b}} = RN, p_{\mathbf{c}} = NR, p_{\mathbf{d}} = NY) \\
&+ \mathbb{P}(p_{\mathbf{a}} = YN, p_{\mathbf{b}} = RN, p_{\mathbf{c}} = NY, p_{\mathbf{d}} = NR) \\
&+ \mathbb{P}(p_{\mathbf{a}} = NR, p_{\mathbf{b}} = NY, p_{\mathbf{c}} = RN, p_{\mathbf{d}} = YN) \\
&+ \mathbb{P}(p_{\mathbf{a}} = NR, p_{\mathbf{b}} = NY, p_{\mathbf{c}} = YN, p_{\mathbf{d}} = RN) \\
&+ \mathbb{P}(p_{\mathbf{a}} = NY, p_{\mathbf{b}} = NR, p_{\mathbf{c}} = RN, p_{\mathbf{d}} = YN) \\
&+ \mathbb{P}(p_{\mathbf{a}} = NY, p_{\mathbf{b}} = NR, p_{\mathbf{c}} = YN, p_{\mathbf{d}} = RN) \qquad (40) \\
&- 4\pi_R\pi_Y\mathbb{P}(p_{\mathbf{a}} = RN, p_{\mathbf{b}} = YN, p_{\mathbf{c}} = NN, p_{\mathbf{d}} = NN) \\
&- 4\pi_R\pi_Y\mathbb{P}(p_{\mathbf{a}} = YN, p_{\mathbf{b}} = RN, p_{\mathbf{c}} = NN, p_{\mathbf{d}} = NN) \\
&- 4\pi_R\pi_Y\mathbb{P}(p_{\mathbf{a}} = NN, p_{\mathbf{b}} = NN, p_{\mathbf{c}} = RN, p_{\mathbf{d}} = YN) \\
&- 4\pi_R\pi_Y\mathbb{P}(p_{\mathbf{a}} = NN, p_{\mathbf{b}} = NN, p_{\mathbf{c}} = YN, p_{\mathbf{d}} = RN) \\
&= 8\pi_R^2\pi_Y^2(1 - e^{-\lambda(l_a+l_b)})(1 - e^{-\lambda(l_c+l_d)}) \\
&- 8\pi_R^2\pi_Y^2(1 - e^{-\lambda(l_a+l_b)}) - 8\pi_R^2\pi_Y^2(1 - e^{-\lambda(l_c+l_d)}) \\
&= 8\pi_R^2\pi_Y^2 e^{-\lambda(l_a+l_b+l_c+l_d)} - 8\pi_R^2\pi_Y^2 \ .
\end{aligned}$$

Similarly,

$$\mathbb{E}\big[w_i(ac|bd)\big|\mathbf{G}_i\big] = 8\pi_R^2\pi_Y^2 e^{-\lambda(l_a+l_b+l_c+l_d+2l_x)} - 8\pi_R^2\pi_Y^2 \ , \qquad (41)$$

and thus,

$$\mathbb{E}\big[w_i(ab|cd) - w_i(ac|bd)\big|\mathbf{G}_i\big] = 8\pi_R^2\pi_Y^2 e^{-\lambda(l_a+l_b+l_c+l_d)}(1 - e^{-2\lambda l_x}). \qquad (42)$$

□

*Proof of Theorem 2.* We prove that the greedy placement algorithm is a statistically consistent estimator of the topology of the true species tree $\mathbf{S}$ by induction:

1. When $\mathbf{S}$ has only four leaves, greedy placement is a statistically consistent estimator.

2. When $\mathbf{S}$ has more than four leaves, let ${S_*}'$ denote the species tree topology of $\mathbf{S}$ without one species $a$. Then, optimally placing $a$ onto ${S_*}'$ is a statistically consistent estimator of the topology of $\mathbf{S}$.

The base case is proven in the proof of Theorem 1; the inductive case can be easily proven by contradiction:

Let $S_*$ be the optimal tree obtained by placing $a$ onto ${S_*}'$. If $S_*$ does not match the topology of $\mathbf{S}$, then $W(S_*) \geq W(\mathbf{S})$, which contradicts Theorem 1. □

| | | | | | | | | | | | | |
|---|---|---|---|---|---|---|---|---|---|---|---|---|
| a | RN | RN | YN | YN | NR | NR | NY | NY | RN | YN | NN | NN |
| b | YN | YN | RN | RN | NY | NY | NR | NR | YN | RN | NN | NN |
| c | NR | NY | NR | NY | RN | YN | RN | YN | NN | NN | RN | YN |
| d | NY | NR | NY | NR | YN | RN | YN | RN | NN | NN | YN | RN |
| w | +1 | | | | | | | | $-4\pi_R\pi_Y$ | | | |
| a | MN | MN | KN | KN | NM | NM | NK | NK | MN | KN | NN | NN |
| b | KN | KN | MN | MN | NK | NK | NM | NM | KN | MN | NN | NN |
| c | NM | NK | NM | NK | MN | KN | MN | KN | NN | NN | MN | KN |
| d | NK | NM | NK | NM | KN | MN | KN | MN | NN | NN | KN | MN |
| w | +1 | | | | | | | | $-4\pi_M\pi_K$ | | | |
| a | WN | WN | SN | SN | NW | NW | NS | NS | WN | SN | NN | NN |
| b | SN | SN | WN | WN | NS | NS | NW | NW | SN | WN | NN | NN |
| c | NW | NS | NW | NS | WN | SN | WN | SN | NN | NN | WN | SN |
| d | NS | NW | NS | NW | SN | WN | SN | WN | NN | NN | SN | WN |
| w | +1 | | | | | | | | $-4\pi_W\pi_S$ | | | |

**Fig. S1**: **Site patterns used in CASTER-pair under the GTR model.** R, Y, M, K, W, S, and N stand for A/G, C/T, A/C, G/T, A/T, C/G, and A/C/G/T, respectively. Here, we define $\pi_R = \pi_A + \pi_G$, ditto for $\pi_Y$, $\pi_M$, $\pi_K$, $\pi_W$, and $\pi_S$. Sites matching multiple patterns are weighted by the total weights of matched patterns.

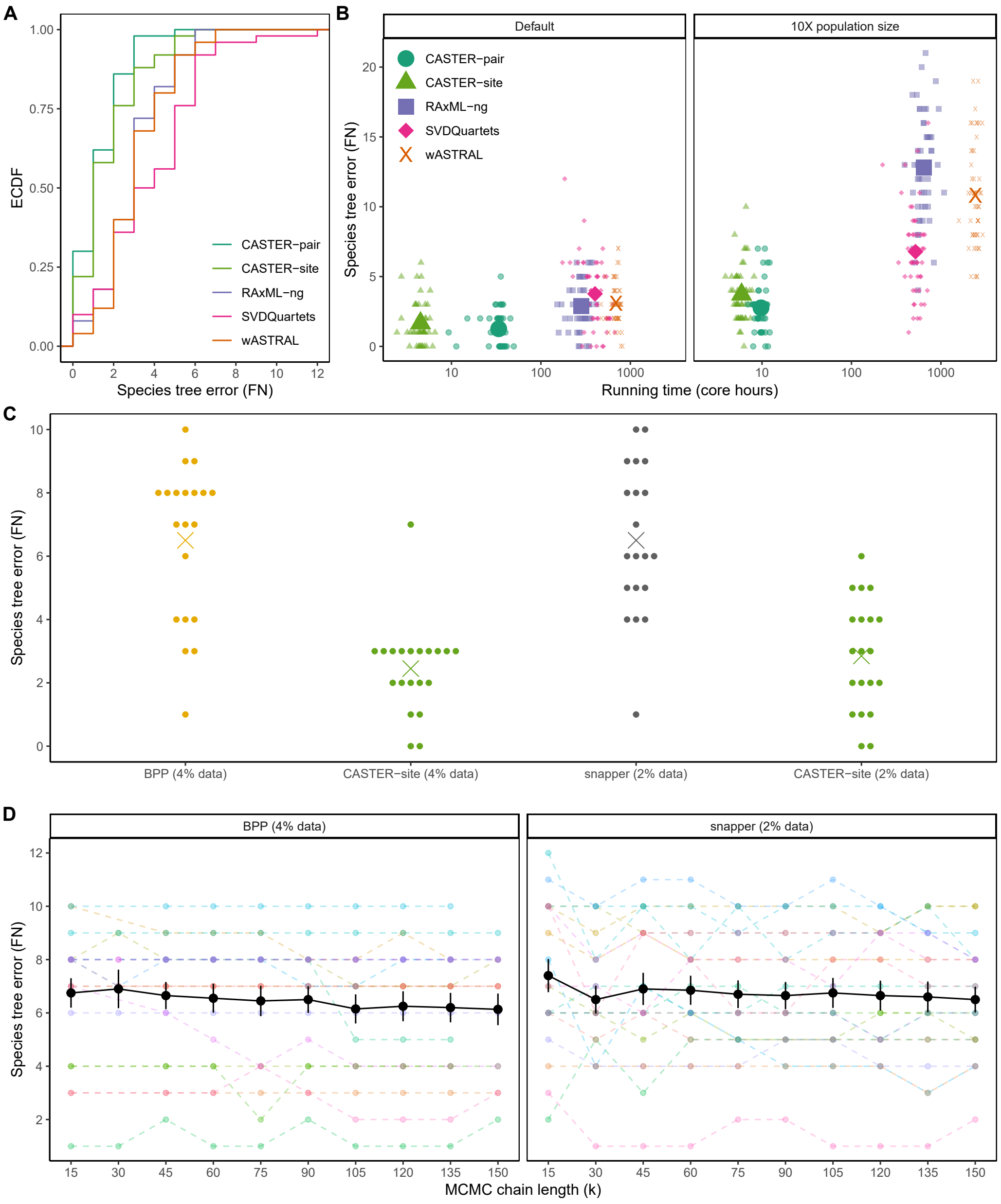


**Fig. S2**: **Additional benchmarking via simulation.** (**A**) Empirical cumulative distribution functions (ECDF) of species tree inference errors of various inference methods in bipartition false negatives (FN) under the default condition of the SR201 dataset. (**B**) Species tree inference errors (FN) versus running time for various methods under 10X population size (compared to the default conditions). (**C**) Species tree inference errors (FN) of BPP and snapper versus CASTER-site with the same input data. (**D**) Species tree inference errors (FN) of consensus trees of BPP and snapper for every 15,000 iterations after a burn-in of 100,000 iterations. Many replicates of BPP fail to finish the entire 150,000 iterations within 25 days (4800 core-hours).

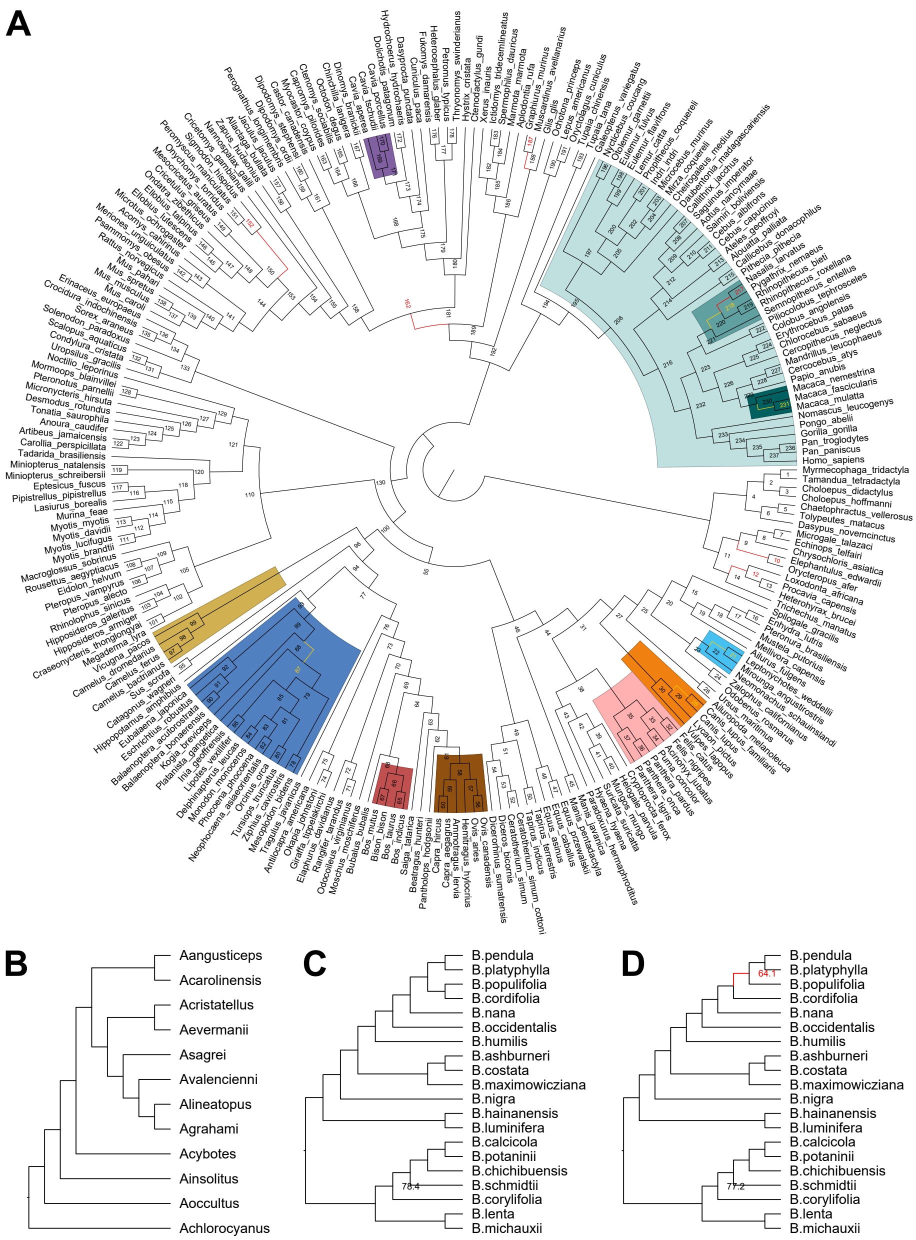


**Fig. S3**: **Inferred species trees by CASTER.** (**A**) CASTER-site tree based on 241 aligned placental mammalian genomes of 1.8 billion informative sites. All branches have 100% bootstrap support. The branches colored red differ from at least one chromosome tree (Fig. 3A), and branches #10 and #217 differ from the published SVDQuartets tree. Clades with abundant introgression signals are highlighted. The branches in Fig. 3E are colored yellow. (**B**) CASTER tree based on 10,390 orthologous genes of 12 lizards. All branches have 100% bootstrap support. (**C**) CASTER-pair and (**D**) CASTER-site tree based on 50,970 loci from 20 diploid birch species. The branches colored red in (B-D) differ from the published ASTRAL/RAxML tree, but match the published ASTRID tree. Branches supports (%) in (B-D) are shown next to the branches, and 100% supports are omitted.

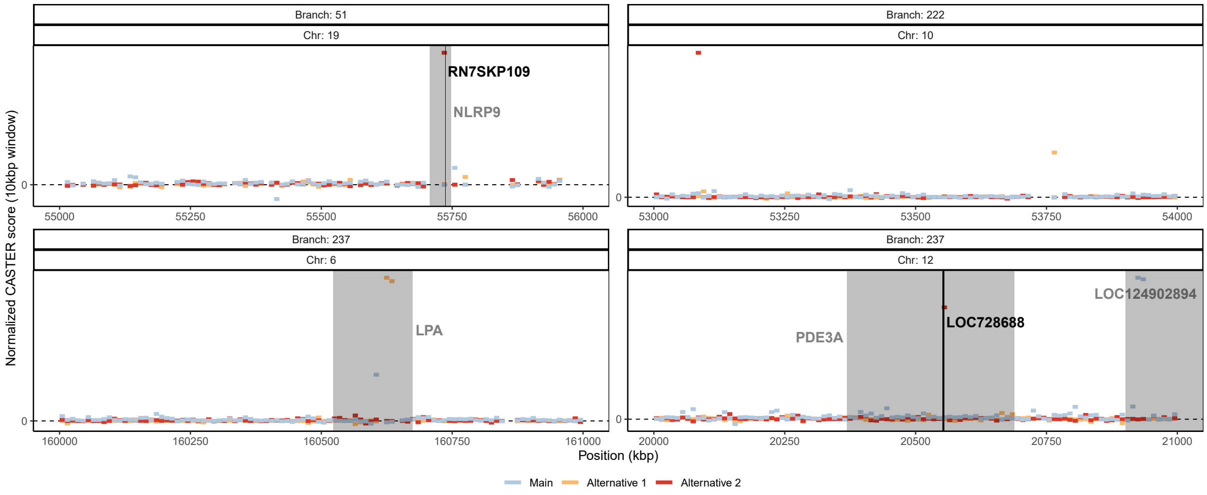


**Fig. S4: Examples of 1kbp windows with abnormally high CASTER scores.** Branch topologies can be found in Fig. S3 using branch numbers. Relevant NCBI gene annotations are highlighted.

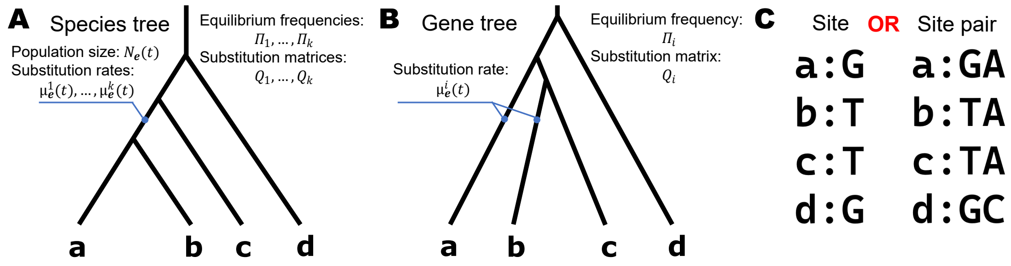


**Fig. S5**: **CASTER assumptions and models.** (**A**) Each species tree branch $\mathbf{e}$, at every moment $t$, has a specific population size $N_{\mathbf{e}}(t)$ and a unique mutation rate $\mu_{\mathbf{e}}^{i}(t)$ for each gene. (**B**) For each gene, the equilibrium frequencies $\Pi_i$ are assumed to be known constants, and the substitution matrix $Q_i$ is also a constant. Each gene tree $\mathbf{G}_i$ is independently sampled under the MSC process, regardless of $\Pi_i$ and $Q_i$. Mutation rates at each moment are passed down from the corresponding species tree branches. (**C**) A single site (or a site pair) is simulated per gene from the gene tree.

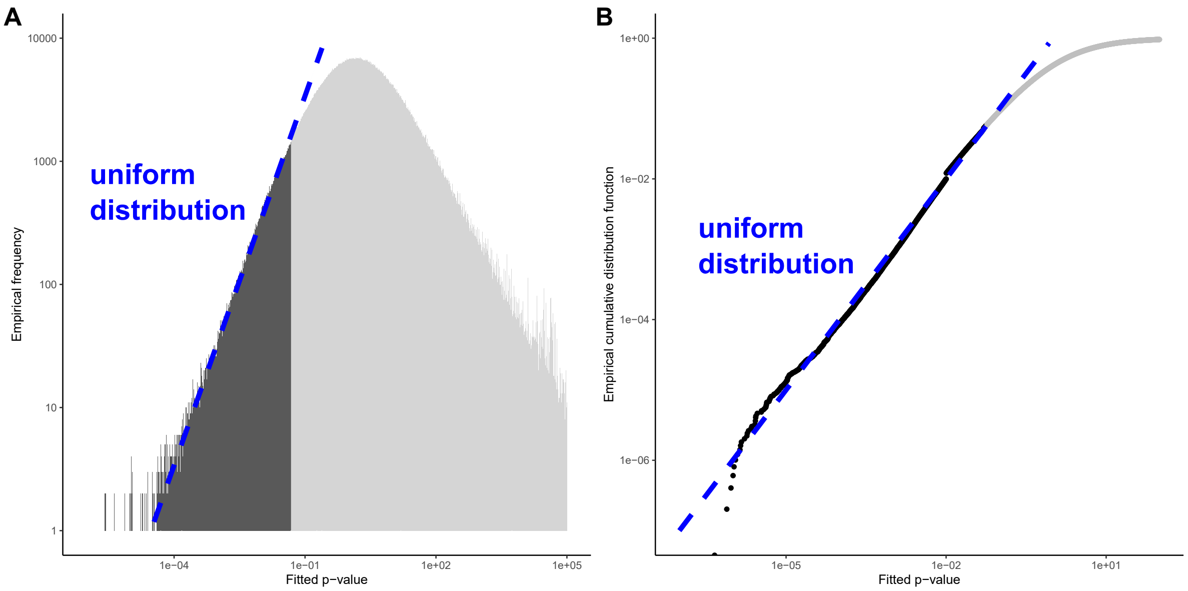


**Fig. S6**: **Empirical design of a null distribution for the normalized CASTER scores.** (**A**) Empirical frequency and (**B**) empirical cumulative distribution function of computed p-values of 10kbp windows on S200 dataset (default condition, 50 replicates, 198 branches, 500 10kbp-windows per branch). The p-values are calculated using the right-hand side of (7). As we are interested in detecting windows with low p-values, we plot both panels in log-log scale and focus only on the p-values $< 0.05$ (high opacity). The blue lines show the uniform distribution between 0 and 1, which is the correct distribution for p-values under the null (note the plots are in log-log scale). Note that deviations from the uniform on the right side are not important as we are only concerned with low p-values (and our claim of exponential approximation was only for low values of $\sigma$). The deviations on the left side are caused by low precision; since $N = 4.95 \times 10^6$, the empirical plots lose precision when $p < 10^{-6}$. Overall, results empirically support the claim that the p-value by equation (7) can be used to test the null hypothesis that a window is generated under the MSC model (ILS only) for any threshold p-value $< 0.05$.

**Table S1**: **P-values for the difference in species tree errors by single-sided paired t-test.** The null hypothesis is that the average accuracy of Condition 1 is no better than Condition 2.

| data | mut | pop | copy | method | Condition 1 | Condition 2 | p-value |
|---|---|---|---|---|---|---|---|
| SR201 | 1X | 1X | 1 | –[a] | CASTER-pair | CASTER-site | 0.005 |
| SR201 | 1X | 1X | 1 | – | CASTER-site | RAxML-ng | $5 \times 10^{-6}$ |
| SR201 | 1X | 10X | 1 | – | CASTER-pair | CASTER-site | $2 \times 10^{-5}$ |
| SR201 | – | 1X | 1 | CASTER-site | 1X | 0.1X | $< 10^{-9}$ |
| SR201 | – | 1X | 1 | CASTER-pair | 1X | 0.1X | $< 10^{-9}$ |
| SR201 | – | 1X | 1 | wASTRAL | 1X | 0.1X | $< 10^{-7}$ |
| SR201 | – | 1X | 1 | RAxML-ng | 1X | 0.1X | 0.006 |
| SR201 | – | 1X | 1 | SVDQuartets | 1X | 0.1X | 0.006 |
| SR201 | 0.1X | 1X | 1 | – | CASTER-pair | RAxML-ng | 0.0001 |
| SR201 | – | 1X | 1 | CASTER-site | 1X | 10X | 0.003 |
| SR201 | – | 1X | 1 | CASTER-pair | 1X | 10X | 0.003 |
| SR201 | – | 1X | 1 | SVDQuartets | 1X | 10X | 0.004 |
| SR201 | – | 1X | 1 | wASTRAL | 1X | 10X | 0.27 |
| SR201 | – | 1X | 1 | RAxML-ng | 1X | 10X | 0.5 |
| SR201 | 10X | 1X | 1 | – | CASTER-pair | RAxML-ng | 0.02 |
| SR201 | 1X | 1X | – | CASTER-site | 2 | 1 | 0.37 |
| SR201 | 1X | 1X | – | SVDQuartets | 2 | 1 | 0.13 |
| SR201 | 1X | 1X | – | RAxML-ng | 2 | 1 | $< 10^{-8}$ |
| SR21 | 1X | 1X | – | CASTER-site | 20 | 5 | 0.72 |
| SR21 | 1X | 1X | – | CASTER-pair | 20 | 5 | 0.009 |
| SR21 | 1X | 1X | – | CASTER-site | 5 | 1 | 0.01 |
| SR21 | 1X | 1X | – | CASTER-pair | 5 | 1 | 0.003 |
| SR21 | 1X | 1X | – | SVDQuartets | 5 | 1 | 0.04 |
| SR21 | 1X | 1X | – | wASTRAL | 5 | 1 | 0.0005 |
| SR21 | 1X | 1X | – | RAxML-ng | 5 | 1 | 0.39 |
| SR21 | 1X | 1X | 2 | – | CASTER-site[b] | BPP | $3 \times 10^{-7}$ |
| SR21 | 1X | 1X | 2 | – | CASTER-site[c] | snapper | $5 \times 10^{-7}$ |

Note: Columns are dataset, mutation rates, effective population size, ploidy/individuals, reconstruction method, testing condition 1, testing condition 2, and p-value by single-sided paired t-test.

[a] A "–" means the values of the column define the conditions being compared.

[b] CASTER-site using 4% data.

[c] CASTER-site using 2% data.

**Table S2**: **Setup and parameters for simulating the default model condition.**

| | | |
|---|---|---|
| Species tree | Demography model | Yule model |
| | Model parameters | $10^6$ generations, $10^{-6}$ speciation rate |
| | # species | 200 + 1 outgroup |
| | Generation multiplier[a] | Gamma$(5, 0.2)$ |
| | Population size[a] | LogNorm$(\ln(10^5) - 0.125, 0.5)$ |
| | # replicates | 50 |
| Chromesome | Ancestry model | Hudson model |
| | Recombination rate | $5 \times 10^{-8}$ per bp per generation |
| | Length | $5 \times 10^6$ bps |
| | # specimens | 1 |
| Sequence | Substitution model | GTR model |
| | Equilibrium frequencies | $(\pi_A, \pi_C, \pi_G, \pi_T) =$ Dirichlet$(36, 26, 28, 32)$ |
| | Substitution matrix | $(r_{AC}, r_{AG}, r_{AT}, r_{CG}, r_{CT}, r_{GT})$ $=$ Dirichlet$(5, 15, 3, 6, 16, 5)$ |
| | Substitution rate | $5 \times 10^{-8}$ |
| | Rate multipliers | Gamma$(\gamma, 1/\gamma)$ shared by Geom$(10^{-4})$ bps and Gamma$(20\gamma, 0.05/\gamma)$ for every bp, where $\gamma =$ LogNorm$(1.5, 1)$ |

Note: Where not specially mentioned, all random samplings are independently performed per replicate.
[a]Those values are drawn independently per species tree branch.

**Table S3**: **The difference in the setup and parameters between the default condition and other model conditions.**

| | | |
|---|---|---|
| Mutation | # replicates[a] | {50,50,20} |
| | Substitution rate | $5 \times 10^{-\{9,8,7\}}$ |
| Population | Population size | LogNorm$(\ln(10^{\{5,6\}}) - 0.125, 0.5)$ |
| Ploidy | # specimens | {1, 2 (diploid unphased)} |
| Scale | # replicates | 1 (first replicate) |
| | Length | $10^6$ bps $\times$ 2048 scaffolds |

Note: Where not specially mentioned, all random samplings are independently performed per replicate.

[a] When using 20 replicates due to computational resource limits, we randomly select replicate number {3,4,11,15,16,22,23,26,27,28,29,31,32,36,38,40,42,43,45,50}.

**Table S4**: **The default setup for each reconstruction method, if not otherwise mentioned.**

| Method | Version | Machine type | # physical cores | # threads |
|---|---|---|---|---|
| CASTER | 1.13.0.0 | AMD EPYC 7742 | 32 | 64 |
| RAxML-ng | 1.1.0 | AMD EPYC 7742 | 64 | 64 |
| SVDQuartets | PAUP* 4.0a | AMD EPYC 7742 | 64 | 128 |
| wASTRAL | 1.13.2.3 | Intel Xeon E5-2670[a] | 1 | 1 |
| | | AMD EPYC 7742 | 16 | 32 |

[a]This setting is used in gene tree reconstructions using RAxML-ng and annotations using IQ-Tree 2.2.0.
